# Three rules for epigenetic inheritance of human Polycomb silencing

**DOI:** 10.1101/2023.02.27.530239

**Authors:** Tiasha A. Shafiq, Danesh Moazed

## Abstract

Heritable expression of developmental genes is mediated by autoregulating transcription factor networks (TFNs) and Polycomb repressive chromatin silencing complexes (PRCs). Whether the mammalian Polycomb pathway mediates DNA sequence-independent epigenetic memory at genes being activated by TFNs remains unknown. Here, we show that, in human cells, newly induced Polycomb silencing at initially active developmental genes can be inherited for many cell divisions. However, when Polycomb silencing is similarly induced near ubiquitously expressed housekeeping genes, it is not heritable. Inheritance requires the recognition of CG-rich DNA by a Polycomb accessory protein, MTF2/PCL2, the histone H3 lysine 27 trimethylation (H3K27me3) and histone H2A lysine 119 mono-ubiquitination (H2AK119ub1) activities of PRC1 and PRC2, and their associated positive feedback (read-write). In the absence of H3K27me3, H2AK119ub1 can mediate inheritance of silencing by a mechanism that requires the recognition of H2AK119ub1 by the variant Polycomb Repressive Complex 1. These findings indicate that inheritance of Polycomb silencing in human cells (1) depends on sequence features of developmental gene loci (2) requires PRC1 and H2AK119ub1, and (3) can be mediated by PRC1-H2AK119ub1 independently of H3K27me3.

## Introduction

Multicellular organisms are composed of many cell types, nearly all of which have the same genome, but express vastly different gene expression programs. Gene expression programs are directed by lineage- and cell type-specific transcription factor networks (TFNs) which form potent autoregulatory loops that must be silenced in other cell types^1–4^. The Polycomb group (PcG) proteins form histone-modifying repressor complexes that silence cell type-specific transcription factors outside their proper spatial domains of expression^5^. Consistent with this important role, mutations in PcG genes are associated with defective gene expression, developmental abnormalities, embryonic lethality, and cancer^6–8^.

Two general models of epigenetic inheritance have been proposed. In the first model, TFNs form trans-acting positive feedback loops that maintain their own expression and concomitantly direct the expression of other cell type-specific genes^4,9,10^. In this model, Polycomb plays a passive role as a default mechanism that silences those genes that are not targeted for activation by cell type-specific TFNs or general transcription factors^11–13^. In the second model, Polycomb plays an active role in inheritance of gene expression programs by forming heritable silent domains that regulate the expression of TFN components and other genes^2,5,13^. Here, positive feedback associated with histone modification and binding activities of Polycomb complexes would maintain the modifications and the silent state during cell division. In support of this model, it has previously been shown that Polycomb silencing induced at a reporter gene inserted in a gene desert can be maintained in the absence of the initiator^14,15^. However, TFNs and general transcription factors are absent in gene deserts, leaving open the possibility that the observed maintenance represents a special case in which Polycomb silencing can self-propagate in the absence of opposing pathways. Whether newly established Polycomb silencing near genes that are controlled by an active TFN and/or general transcription factors can withstand erasure and be inherited in the absence of recruitment signals remains unknown.

The PcG genes were identified based on mutations that give rise to homeotic transformations in *Drosophila*^16–18^. Subsequent studies showed that PcG proteins are highly conserved and form multiple Polycomb repressive complexes (PRCs) that have histone modifying and binding activities^5^. In mammals, canonical PRC1 (cPRC1) is composed of a heterodimer of PCGF2 or 4 with the RING1A or RING1B E3 ubiquitin ligases, which mono-ubiquitinate histone H2AK119, and two additional subunits, CBX2, 4, 6, 7, or 8 chromobox proteins and PHC1, 2, or 3^19–21^. The PRC2 complex contains EED, SUZ12, RBBP7 or RBBP8, and the EZH1 or EZH2 H3K27 methyltransferases^22–25^. In addition, variant PRC1 (vPRC1) complexes have been identified that lack CBX subunits and contain RYBP or YAF2 and one of four PCGF proteins (PCGF1, 3, 5, or 6)^26^.

Polycomb complexes contain both catalytic (writer) and substrate recognition (reader) subunits that mediate extensive cross talk between them^27–29^. The EED subunit of PRC2, with its seven WD40 repeat domains, binds H3K27me3 and allosterically activates the EZH2 methyltransferase^27,30–32^. H3K27me3 is also recognized by the CBX subunits of PRC1, which mediate chromatin compaction and silencing^33–35^. H2AK119ub1 is recognized by the RYBP/YAF2 subunits of vPRC1^36–38^ in addition to the JARID2 and AEBP2 accessory subunits of PRC2^39,40^. H2AK119ub1 plays a key role in mammalian Polycomb silencing. Deletion of RING1A and RING1B resulting in the loss of H2K119ub1 leads to the concomitant loss of most H3K27me3 domains in mouse embryonic stem cells^41,42^. In addition the two modification systems perform mutually dependent as well as independent silencing functions in mouse zygotes and early embryos^43,44^. Despite its critical role, it remains unclear whether H2AK119ub1-mediated silencing in the absence of H3K27me3, observed in mESCs^41,42,45^, early mouse embryos^43,44^, and human HEK293 cells^46^, represents an epigenetically heritable silencing modification.

In addition to forming interwoven histone modification and binding activities, PRC1 and PRC2 interact with accessory factors that bind to DNA. The mammalian vPRC1/PCGF1 associates with the KDM2B Jumonji domain (JmjC)-containing protein, which binds to unmethylated CpG islands (CGIs) through its CXXC zinc finger domain^47^. The PRC2 complex interacts with multiple DNA-binding proteins, including JARID2 and Polycomb-like proteins (PCL), PCL1/PHF1, PCL2/MTF2, and PCL3/PHF19, which bind to CGIs using their winged helix motif in the extended homology domain^5,48–50^. Deletion of individual PCL protein, or mutations that disrupt the ability of PCL2/MTF2 to bind DNA, result in reduced PRC2 recruitment and H3K27me3 at most Polycomb target genes suggesting an important role for CGI recognition^48–50^. However, composite DNA elements that bind to multiple transcription factors, akin to *Drosophila* Polycomb response elements (PREs), are thought to be absent in mammals^5^. Furthermore, since many Polycomb repressed genes are transcriptional regulators that form components of cell type-specific TFNs, Polycomb repression must contend with the potential autoactivation of these factors to prevent their ectopic expression. Whether Polycomb can act in a dominant manner to forestall TFN activation and whether specific features of developmental genes, such as CGIs, are required for the inheritance of Polycomb domains remain unknown.

In this study, we used an inducible reporter gene silencing system to show that Polycomb silencing could be established near or at both ubiquitously expressed housekeeping genes and developmental genes (which have active TFNs in the cell type used) but is only heritable at the latter loci. Inheritance requires the ability of PCL2/MTF2 to recognize CGIs, the EED and SUZ12 subunits of PRC2, and the RING1A/B H2AK119 E3 ubiquitin ligase subunit of PRC1. However, aromatic cage mutations that abolish the ability of EED to recognize H3K27me3 and lead to loss of H3K27me3 weaken but do not abolish inheritance. Notably, in the absence of detectable H3K27me3, mutations that impair the ability of RYBP to recognize H2AK119ub1 lead to a complete loss of heritable silencing. These findings suggest that, in addition to read-write positive feedback loops, Polycomb memory requires the recognition of specific DNA-sequences at developmental genes. In addition, they identify a primary role for H2AK119ub1 in epigenetic inheritance of Polycomb silencing.

## Results

### Locus-dependent inheritance of Polycomb silencing

To investigate the role of DNA context on inheritance of Polycomb silencing, we induced Polycomb silencing at two types of genes: developmentally regulated genes that are targeted for silencing by Polycomb in some cell types but are active in human embryonic kidney cells (HEK293FT) and genes that are not targeted by Polycomb, such as ubiquitously expressed housekeeping genes, which are active in HEK293 and other cell types. We integrated a reporter cassette with 5 tetracycline operators (*5xtetO*) upstream of the *EF1* promoter driving *H2B-CITRINE* expression (*5xtetO-H2B-CITRINE*) a few kilobases upstream of multiple Polycomb and non-Polycomb target loci in HEK293FT cells (**Figure 1A**). In the same cell lines, we also expressed a reverse Tet repressor (rTetR, Tet-ON) protein fused to the CBX7 subunit of cPRC1 to initiate silencing (**Figure 1A**). The rTetR-CBX7 protein only binds the *5xtetO* array in the presence of doxycycline (+Dox) and is released upon removal of doxycycline (Dox Removal) from the culture medium, allowing us to control the association of the initiator with DNA and assess silencing, H3K27me3, and H2AK119ub1 with and without DNA sequence-dependent initiation (**Figure 1A**).

**Figure 1.**
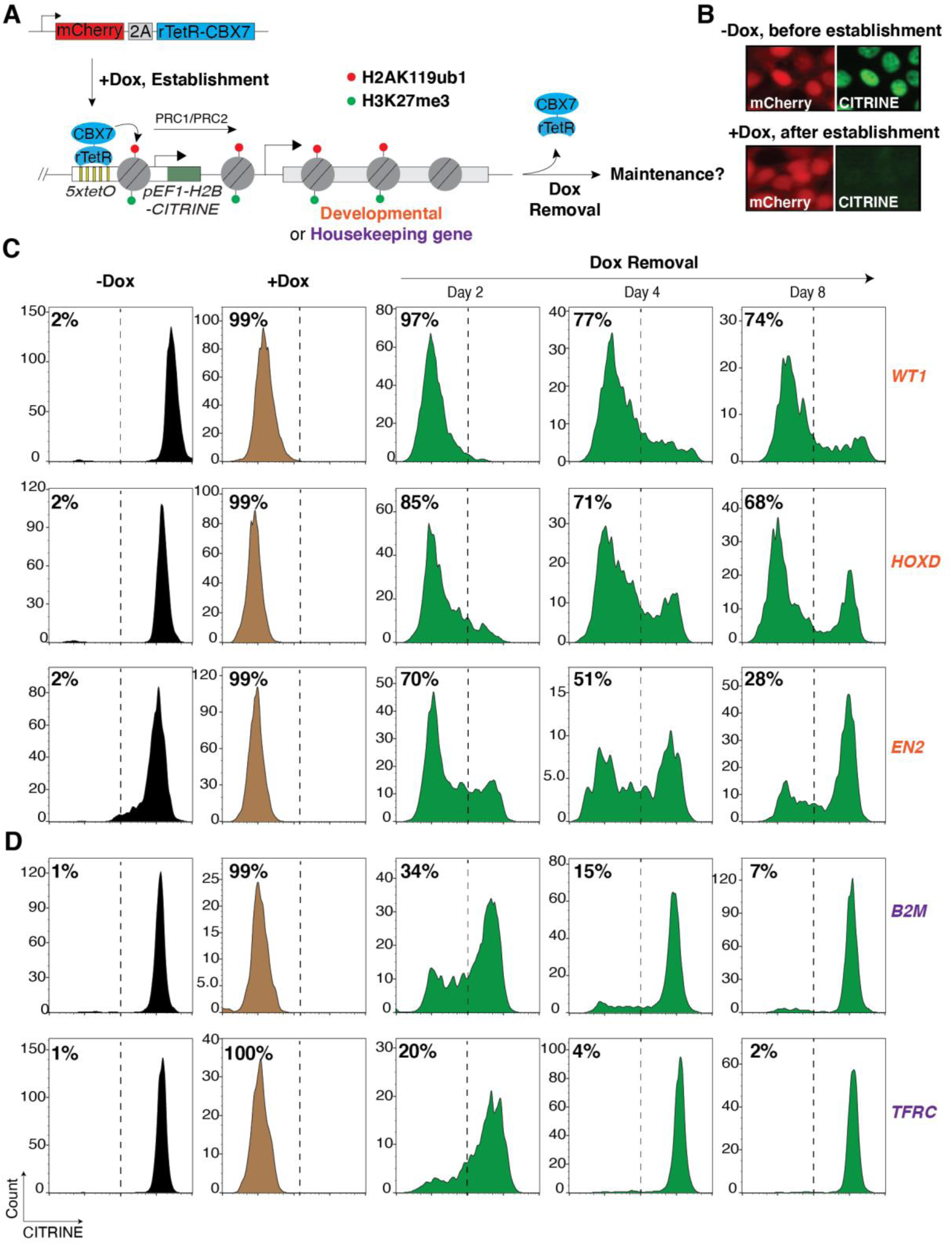
Newly established Polycomb silencing near developmental genes, but not at ubiquitously expressed genes, is heritable. **A**, Schematic diagram illustrating the strategy for inducible Polycomb silencing by expression of rTetR-CBX7 as a fusion with mCherry-2A in cells carrying the *5xtetO-CITRINE* reporter at various developmental and ubiquitously expressed genes. See Figures S1 and S2 for exact coordinates of reporter insertion. **B**, Representative fluorescence images showing mCherry and CITRINE expression before and after establishment of silencing (-Dox and +Dox, respectively). **C**, Flow cytometry histograms showing CITRINE expression before and after establishment of silencing with doxycycline addition (-Dox and +Dox) and at the indicated days after removal of doxycycline (Dox removal) at developmental genes *WT1*, *HOXD* and *EN2*. Percentages (%) indicate the fraction of CITRINE negative cells. **D**, Same as **C**, but with the reporter inserted near ubiquitously expressed genes *B2M* and *TFRC*.

At the Polycomb target loci *WT1*, *EN2*, *HOXD and HOXB*, which encode developmentally regulated transcription factors (Wilms tumor, Engrailed 2, and homeobox transcription factors, respectively), we inserted the *5xtetO-H2B-CITRINE* reporter 3-4.5 kb upstream of the endogenous promoter regions (**Figure 1A, Figure S1A-D**). In HEK293FT cells, these genes are devoid of H3K27me3 and are expressed, while in human embryonic stem cells (hESCs), they are associated with H3K27me3 and are not expressed (**Figure S1A-D**). We cultured cells in +Dox medium to establish silencing for 8 days (∼8 cell divisions), with cells in doxycycline-free (-Dox) medium serving as controls (**Figure 1B**). Fluorescent activated cell sorting (FACS) analysis showed that the *CITRINE* reporter was silenced in >95% of cells, in cell lines with the reporter inserted at the *WT1*, *EN2*, and *HOXD* loci, grown in +Dox medium but was fully expressed in cells grown in -Dox medium in which rTetR-CBX7 does not bind the *5xtetO* sites (**Figure 1C**). The *CITRINE* reporter inserted at the *HOXB* locus was poorly silenced and was not further pursued (**Figure S1E**). Following establishment, doxycycline was removed to release the rTetR-CBX7 from the *5xtetO* site (Dox Removal). As shown in **Figure 1C**, silencing of the *CITRINE* reporters at the *WT1* and *HOXD* loci was maintained at 2, 4 and 8 days after release of the rTetR-CBX7 initiator, with >70% of the cells maintaining silencing at day 8 at the *WT1* locus. Silencing was also maintained at the *EN2* locus but displayed more rapid decay kinetics (**Figure 1C**). At the *WT1* locus, 40 days after release of rTetR-CBX7, 27% of the cells still maintained silencing (**Figure S1F**), and we further verified the inheritance of silencing by CITRINE fluorescence imaging at 2, 4, and 8 days after Dox removal (**Figure S1G**). These results indicate that transient rTetR-CBX7 recruitment near developmental genes leads to heritable silencing.

As controls, tethering rTetR alone did not lead to silencing of the *CITRINE* reporter, indicating that silencing was not caused by rTetR-mediated steric inhibition (**Figure S1H**). Moreover, silencing was maintained when we deleted rTetR-CBX7 8 days after the establishment of silencing, demonstrating that leaky binding of rTetR-CBX7 was not responsible for epigenetic inheritance of silencing at the *WT1* locus (**Figure S1I**).

We next inserted the *5xtetO-H2B-CITRINE* reporter at the endogenous promoter regions of two ubiquitously expressed housekeeping genes: 4.7 kb upstream of Beta-2-Microglobulin, *B2M*, or 1 kb upstream of the transferrin receptor, *TFRC*. Ubiquitously expressed genes are regulated by general transcription factors, are expressed in all cell types, and are devoid of Polycomb modifications^51^. In HEK293FT cells, H3K27me3 and H2AK119ub1 are absent at the *B2M* and *TFRC* genes (**Figure S2A, B**). We carried out establishment and maintenance assays as we did for the Polycomb target genes. Silencing was robustly established at both loci as the *5xtetO-H2B-CITRINE* reporter was silenced in >95% of cells (**Figure 1D**). However, in contrast to the developmentally regulated genes (**Figure 1C**), reporter gene silencing was rapidly lost at these loci (**Figure 1D**). While about 34% (*B2M*) or 20% (*TFRC*) of the cells maintained silencing 2 days after the release of rTetR-CBX7, only 15% (B2M) or 4% (TFRC) and 7% (*B2M*) or 2% (*TFRC*) maintained silencing by 4 and 8 days after release, respectively (**Figure 1D, Figure S2C**). These results indicate that Polycomb-mediated silencing established near ubiquitously expressed genes, *TFRC* and *B2M,* was not epigenetically inherited.

### Heritable silencing is associated with inheritance of Polycomb modifications at the reporter gene and adjacent developmental gene

We next examined the establishment and maintenance of Polycomb-associated histone modifications at the developmental *WT1* and housekeeping *B2M* and *TFRC* loci. At the *WT1* locus, ChIP-qPCR and ChIP-seq results showed that a domain of H3K27me3 and H2AK119ub1 was established by rTetR-CBX7 and, consistent with maintenance of reporter gene silencing, these modifications were maintained 8 days after the release of rTetR-CBX7 (**Figure 2A-C, Figure S3A**). The H3K27me3 domain extended to ∼7 kb on both sides of the *tetO* sites and spread to the promoter and first exon of the *WT1* gene and was enriched at levels comparable to an endogenous Polycomb target gene, *MYT1* (**Figure 2A-B**). Importantly, the newly established H3K27me3 and H2AK119ub1 spanning endogenous *WT1* sequences were maintained 8 days after of the release of rTetR-CBX7 (**Figure 2A-C, Figure S3A**, highlighted in red), indicating that the establishment and epigenetic inheritance of the Polycomb domain was not restricted to the reporter gene. On the other hand, at the *B2M* and *TFRC* housekeeping loci, H3K27me3 and H2AK119ub1 were enriched at regions surrounding the reporter cassette but were lost in the maintenance phase, 8 days after the release of rTetR-CBX7 (**Figure 2D-E; Figure S3B-E**). In contrast to the *WT1* locus, H3K27me3 and H2AK119ub1 did not spread to the promoter region of either *B2M* or *TFRC* genes (**Figure 2D-E; Figure S3B-E**). These results demonstrate that the establishment of silencing at both the developmental and housekeeping genes is coupled to histone H3K27me3 and H2AK119ub1, which can be epigenetically maintained for several cell divisions after release of the rTetR-CBX7 initiator only at developmental genes.

**Figure 2.**
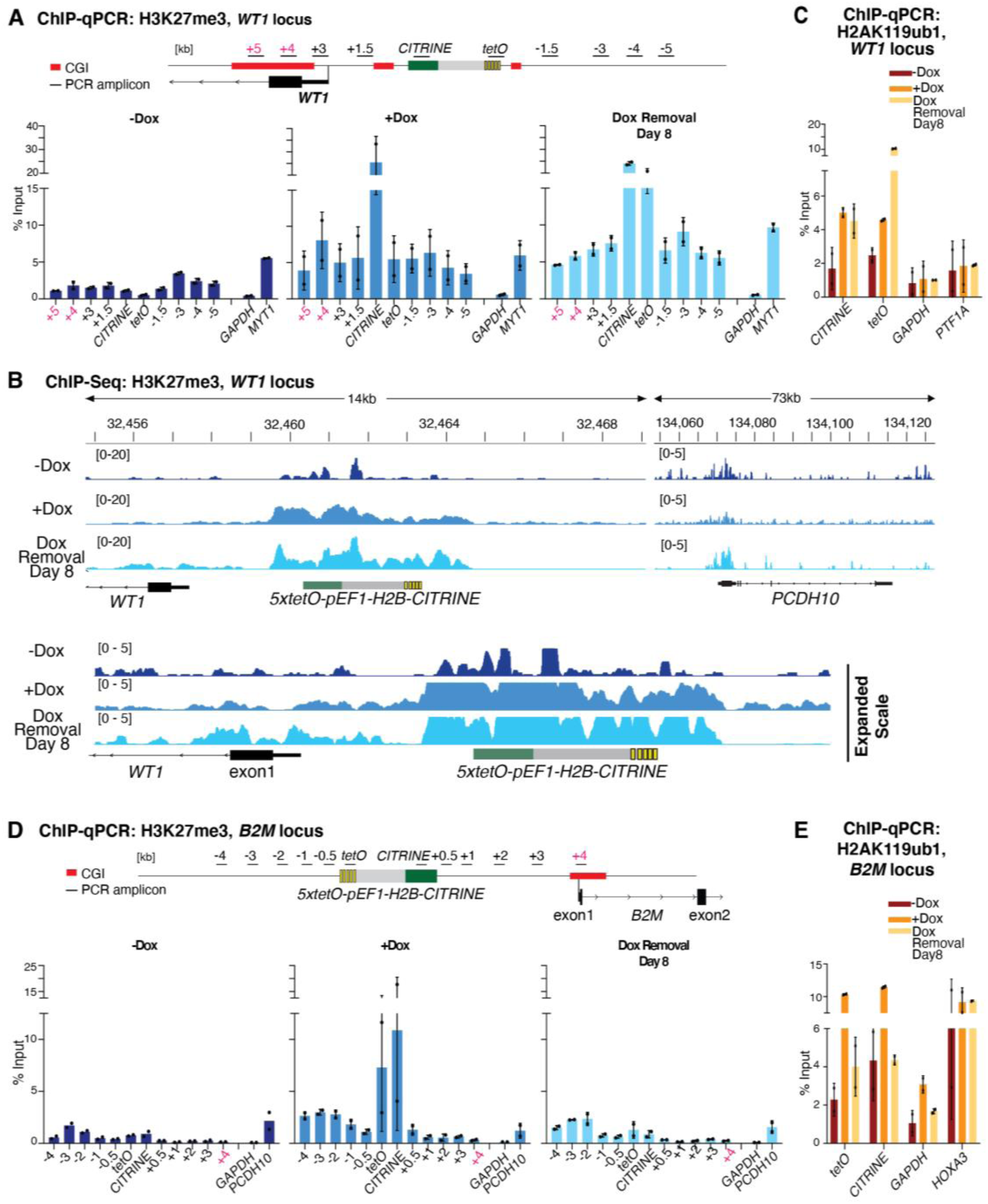
Memory of newly established H3K27me3 and H2AK119ub1 near or at the developmental gene, *WT1*, but not near the ubiquitously expressed gene, *B2M*. **A,** ChIP-qPCR analysis of H3K27me3 enrichment at the reporter locus and surrounding regions at the *WT1* locus before establishment (-Dox), after establishment (+Dox), and during the maintenance phase (Dox removal). *GAPDH* served as a negative control and the native Polycomb-repressed *MYT1* gene as a positive control. Results for 2 biological replicates are presented. Error bars represent standard deviations. The location of PCR amplicons and CGIs are highlighted in the map at the top. **B**, Genome browser snapshots of H3K27me3 ChIP-seq reads at the reporter locus inserted near *WT1* and the endogenous Polycomb-silenced *PCDH10* genes. Expanded scale (bottom) highlights the spreading of H3K27me3 to the promoter of WT1 and its maintenance. **C,** Same as **A** but H2AK119ub1 ChIP-qPCR. **D**, ChIP-qPCR analysis of H3K27me3 enrichment at the reporter locus inserted near *B2M* and surrounding regions before establishment (-Dox), after establishment (+Dox), and during the epigenetic maintenance of silencing (Dox removal). *GAPDH* served as a negative control and the native Polycomb-repressed *PCDH10* gene as a positive control. Results for 2 biological replicates are presented. Error bars represent standard deviations**. E,** Same as **D** but H2AK119ub1 ChIP-qPCR; *HOXA3* served as a positive control.

To rule out the effect of differences in rTetR-CBX7 expression levels in inheritance of silencing at the developmentally regulated genes versus ubiquitously expressed genes, we used western blotting to show that rTetR-CBX7 was expressed at similar levels in cell lines with the reporter inserted at different loci (**Figure S4A**). Furthermore, to verify rTetR-CBX7 binding and release, we constructed rTetR-CBX7-Flag cell lines, which expressed similar levels of rTetR-CBX7-Flag and displayed heritable silencing at the *WT1* but not the *TFRC* locus (**Figure S4B, C**). Experiments using these cell lines showed that rTetR-CBX7 localized with similar efficiency to the reporter gene at the *WT1* and *TFRC* loci under establishment conditions (**Figure S4D**). At the *TFRC* reporter, no rTetR-CBX7 binding was detected 4 days after its release by growth in medium lacking Dox. Interestingly, at the *WT1* locus, there was binding of the rTetR-CBX7 under maintenance conditions, albeit to lower levels than in the establishment phase. This is likely due to the incorporation of rTetR-CBX7 into the PRC1 complex and its recruitment to the locus during maintenance phase via Polycomb histone modifications, independently of the *5xtetO* array. As expected, in cells grown in -Dox medium, rTetR-CBX7 did not bind to the reporter locus, as the ChIP-qPCR signals were similar to the cell line where rTetR-CBX7 was absent (**Figure S4D**). This result further confirmed that there was no leaky binding of rTetR-CBX7 to the 5x*tetO* array in the absence of Dox.

To test whether proximal transcriptional activity at ubiquitously expressed genes blocks the inheritance of silencing at the *5xtetO-H2B-CITRINE* reporter, we deleted specific sequences at the promoter regions of these gene. We deleted a 4.4 kb region between the *CITRINE* reporter and the end of exon 1 of the *B2M* gene (*B2MΔ1*)(**Figure 3A, Figure S5A-B**). This deletion removed the majority of c-Myc (a general transcription factor) binding sites and dramatically reduced *B2M* mRNA and protein levels (**Figure 3B; Figure S5A-C**). Two smaller homozygous sequence deletions at the *B2M* locus (*B2MΔ2 and B2MΔ3*) similarly reduced mRNA levels (**Figure 3B, Figure S5D-K**). However, while *B2MΔ1* cells displayed weak short-term inheritance of *CITRINE* reporter silencing, the *B2MΔ2 and B2MΔ3* cells did not result in heritable silencing (**Figure 3C; Figure S5K**), suggesting that the lack of inheritance was not simply due to high levels of transcription at the *B2M* locus. At the *TFRC* locus, we deleted a 5.6 kb region between the reporter and the end of exon 2 of the *TFRC* gene (*TFRCΔ1*)(**Figure 3D, Figure S6A-B**). This deletion eliminates binding sites for c-Myc, a general transcriptional factor that enhances *TFRC* expression^52,53^, but is also likely to delete other binding sites (**Figure 3D; Figure S6A-B**). As shown by qRT-PCR analysis of RNA levels, steady state mRNA and proteins levels were greatly reduced in *TFRCΔ1* cells (**Figure 3E**, **Figure S6C).** The *TFRCΔ1* cells showed a longer delay in the loss of silencing with 53% and 35% of the cells silenced at Day 4 and Day 8 after rTetR-CBX7 release, respectively, relative to 4% in control cells (**Figure 3F**). Furthermore, ChIP-qPCR showed the presence of H3K27me3 during maintenance at 4 days after Dox removal (**Figure S6D**). A shorter 0.8 kb homozygous sequence deletion, *TFRCΔ2*, showed only a ∼2-fold reduction in steady state mRNA levels, exhibited only a modest delay in the loss of maintenance of reporter (**Figure 3D-F, Figure S6E-H)**. Altogether, these results indicate that DNA deletions that reduce transcription allow short-term inheritance of reporter gene silencing near housekeeping genes, and suggest that long-term epigenetic inheritance is associated with specific sequence features of developmental genes.

**Figure 3.**
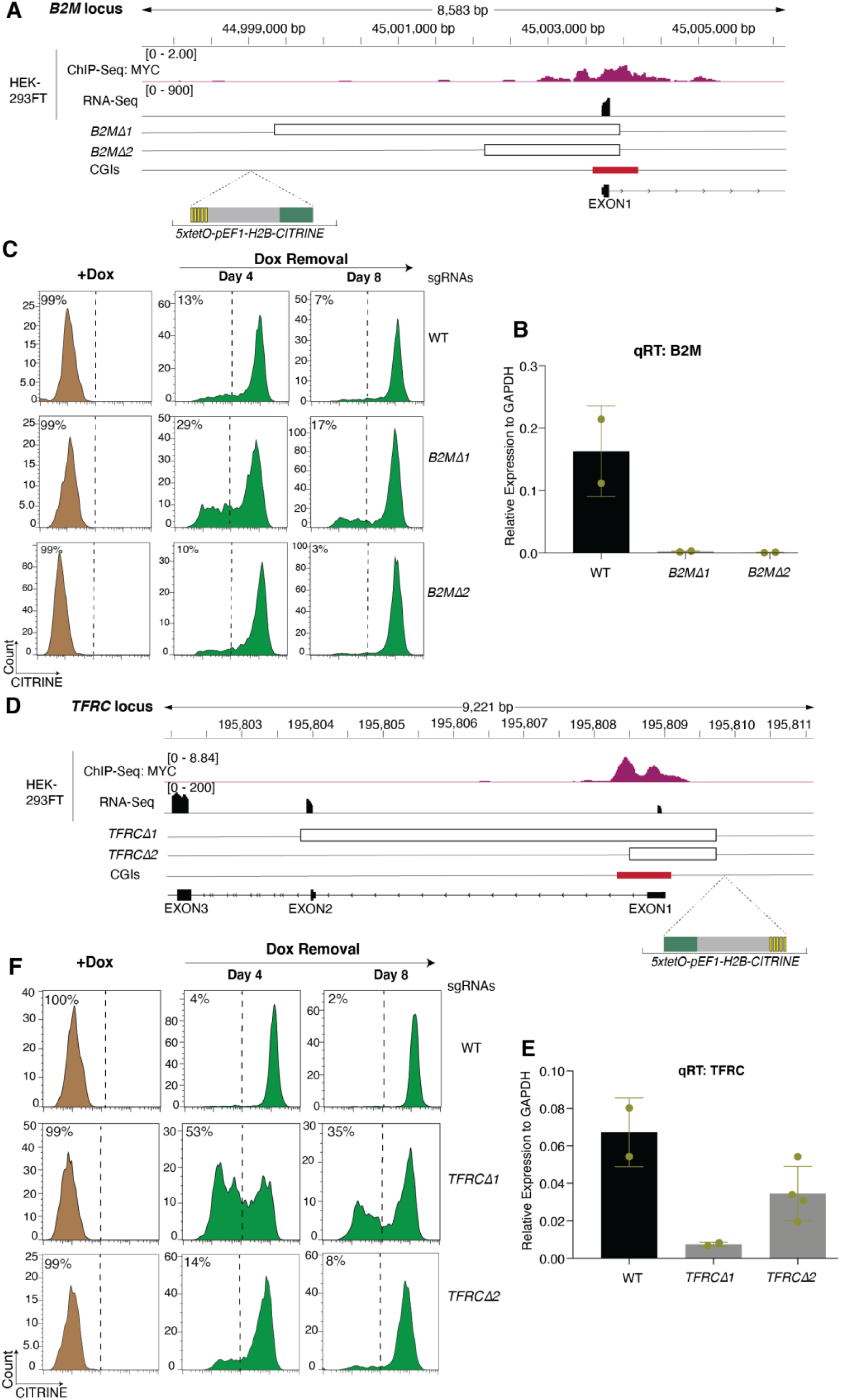
*Cis* proximal transcriptional activity is not a major determinant for epigenetic inheritance of Polycomb silencing at ubiquitously expressed genes. **A,** Genome browser tracks showing publicly available chromatin features of the *B2M* locus in HEK293FT cells (see Table S8 for references). Chromosome coordinates (top), c-Myc transcription factor binding sites, RNA-seq reads, deleted promoter regions DNA (*B2M Δ1* and *B2M Δ2*), and reporter insertion site (bottom) are indicated. **B,** qRT-PCR plot detecting steady state B2M mRNA levels in wild-type control and *B2M Δ1* and *B2M Δ2* cell lines shown in **A**. B2M RNA levels were normalized to GAPDH RNA levels. Error bars represent standard deviations. **C**, Flow cytometry histograms showing CITRINE expression after establishment (+Dox) and at the indicated days after removal of doxycycline (Dox removal) in wild-type control and *B2M Δ1* and *B2M Δ2* cell lines with deletions of endogenous promoter sequences at *B2M* locus. Percentages (%) indicate the fraction of CITRINE negative cells. **D,** Same as **A** but for endogenous promoter deletions at the *TFRC* locus. **E**, same as **B**, but showing RNA levels in *TFRCΔ1* and *TFRCΔ2* cells. **F**, same as **C**, but showing flow cytometry histograms for *TFRCΔ1* and *TFRCΔ2* cells.

### Recognition of CG-rich DNA by MTF2-PRC2 is required for epigenetic inheritance

The core PRC2 complex associates with accessory subunits that bind CG-rich DNA at Polycomb target genes^48–50^, including association with PCL1-3 proteins to form PRC2.1 and with JARID2 to form PRC2.2^54–56^. Additionally, developmental genes that are targeted for Polycomb silencing have been noted to contain a higher density of CpG islands (CGIs)^48,57,58^ and the developmental genes studies here all had more annotated CGIs that extended beyond the promoter-associated CGI present at the ubiquitously expressed genes (**Figure S7A**). We used knockout and rescue experiments to investigate the possible role of the above DNA binding proteins and their contact with CGIs at developmentally regulated genes in inheritance of Polycomb mediated gene silencing. Deletion of *PCL2/MTF2* (*MTF2^-/-^)* showed strong loss of *CITRINE* reporter silencing at the *WT1* locus by 8 days after release of the rTetR-CBX7 and near complete loss by 16 days after release (**Figure 4A, Figure S7B**), indicating that MTF2 was required to maintain silencing at the *WT1* locus. MTF2 binds to CG-rich DNA through its Extended Homology Domain^48,49^. To determine whether the DNA binding activity of MTF2 was necessary for the maintenance of the *CITRINE* reporter silencing, we introduced 3x-Flag-tagged wild-type MTF2 (Flag-MTF2) or mutant Extended Homology Domain MTF2 (K338A, K339A; referred to as Flag-MTF2-EH), which does not bind DNA in vitro, into *MTF2^-/-^* cells (**Figure 4B, Figure S7C**). In *MTF2^-/-^* cells overexpressing wild-type Flag-MTF2, maintenance of *CITRINE* reporter silencing was restored, while in *MTF2^-/-^* cells overexpressing the Flag-MTF2-EH mutant maintenance was not restored (**Figure 4C**). ChIP-qPCR and ChIP-Seq for *Flag-MTF2* rescue cells demonstrated that wild-type Flag-MTF2 was recruited during establishment and maintenance (**Figure 4D, Figure S7D**). By contrast, Flag-MTF2-EH was not recruited during establishment or maintenance (**Figure 4D, Figure S7D**), suggesting that the interaction of MTF2 with DNA was required for its efficient recruitment. Furthermore, wild-type Flag-MTF2 but not Flag-MTF2-EH was enriched at the endogenous *FOXQ1* Polycomb target gene (**Figure S7D**). ChIP-qPCR and ChIP-Seq in *MTF2^-/-^* showed reduced H3K27me3 levels during maintenance at the *WT1 CITRINE* reporter, which were restored in wild-type *Flag-MTF2* cells during maintenance (**Figure 4E, Figure S7E**). However, in mutant *Flag-MTF2-EH* cells, H3K27me3 levels were not restored, and were even reduced to a greater extent, likely due to a dominant negative effect of the mutant (**Figure 4E, Figure S7E**). As expected, *Flag-MTF2-EH* cells had reduced H3K27me3 levels at the endogenous Polycomb target genes *MYT1* and *PCDH10* (**Figure 4E, Figure S7E**). These results demonstrate that the DNA binding activity of MTF2 is required for its efficient recruitment by rTetR-CBX7 and for maintenance of Polycomb silencing at the *WT1* locus.

**Figure 4.**
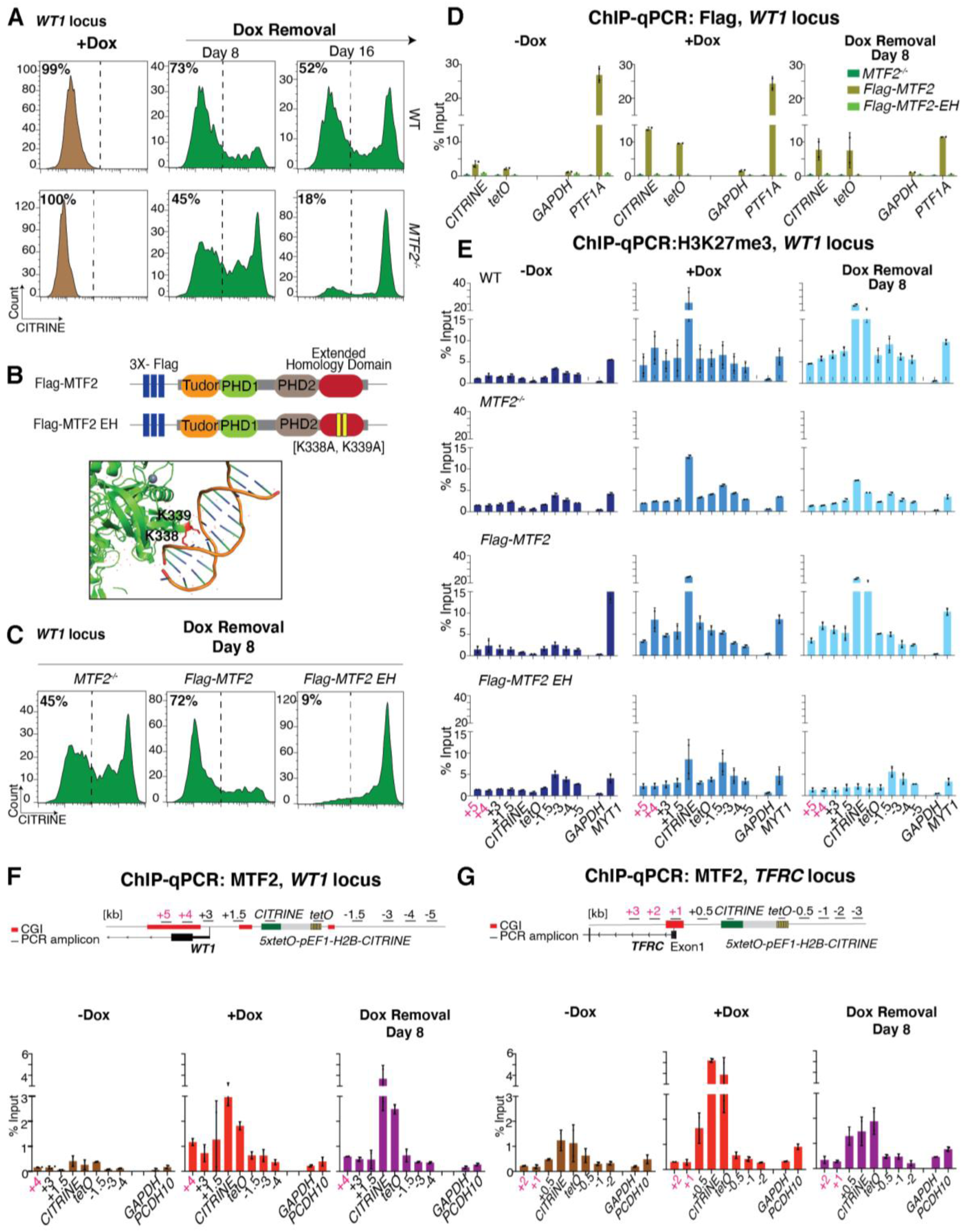
The DNA binding activity of *MTF2* is required for epigenetic inheritance of Polycomb silencing at the *WT1* developmental gene. **A**, Flow cytometry histograms showing CITRINE expression, inserted at the Polycomb target locus*, WT1*, after establishment of silencing (+Dox) and at the indicated days after removal of doxycycline (Dox removal) in wild-type control (WT) and *MTF2* deleted cell line (*MTF2^-/-^)*. **B**, Schematic diagram showing the wild-type MTF2 (*Flag-MTF2*) and the two residue mutations [K338A, K339A] generated in the extended homology (EH) domain of MTF2 (*Flag-MTF2-EH*) (top). A snapshot of the structure of *MTF2* EH domain with bound DNA. MTF2 is colored in green, the K338 and K339 residues are in red, and the DNA is multicolored. (PDB5XFR)^49^. **C**, Flow cytometry histograms showing CITRINE expression at Day 8 after removal of doxycycline (Dox removal) in *MTF2^-/-^* and *MTF2^-/-^* cells overexpressing Flag-MTF2 or Flag-MTF2-EH. **D**, ChIP-qPCR analysis of Flag enrichment at the reporter locus and surrounding regions before establishment (-Dox), after establishment (+Dox), and 8 days after removal of doxycycline (Dox removal) in *MTF2^-/-^* and *MTF2^-/-^* overexpressing Flag-MTF2 and Flag-MTF2-EH. *GAPDH* served as a negative control and the native Polycomb-repressed *PTF1A* gene as a positive control. Error bars represent standard deviations. **E,** Same as **D**, but ChIP for H3K27me3 in WT, *MTF2^-/-^* and *MTF2^-/-^* overexpressing Flag-MTF2 or Flag-MTF2-EH. **F-G,** same as **D**, but ChIP for MTF2 at the *WT1* **(F)** and *TFRC* **(G)** loci.

To test whether other PRC2 accessory proteins affect inheritance, we deleted three other DNA-binding proteins that are accessory to PRC2, PRC2.2 subunit JARID2 and PRC2.1 subunits PHF1 (PCL1) and PHF19 (PCL3) in the cell line containing the *CITRINE* reporter at the *WT1* locus. We found that deletion of *JARID2* and *PHF19* had no effect on establishment or maintenance (**Figure S8A, B**), but the deletion of *PHF1* led to slow loss in maintenance by Day 16 after rTetR-CBX7 release, similar to *MTF2^-/-^* cells (**Figure S8C**). Therefore, MTF2 and PHF1 may compensate for one another to a limited extent, but epigenetic maintenance is eventually lost in the absence of either factor. Together, these results suggest that the CGI binding ability of PRC2 accessory factors MTF2 and PHF1 is important for epigenetic inheritance of silencing at the *WT1* locus.

To further investigate whether the inheritance of Polycomb silencing correlated with MTF2 recruitment, we used a commercially available antibody to compare MTF2 localization at the *TFRC* and *WT1* loci. ChIP-qPCR experiments showed that MTF2 was recruited to the reporter gene at both loci during establishment of silencing in +Dox medium and was efficiently maintained at the *WT1* locus during the maintenance phase 8 days after Dox removal (**Figure 4F, G**). Importantly, during the establishment phase (+Dox), MTF2 was enriched at the reporter as well at CGIs extending 4 kb downstream of the *WT1* promoter; by contrast, at the *TFRC* locus, MTF2 enrichment was more restricted and did not spread to the lone promoter-associated CGI (**Figure 4F, G**). This suggests that additional non-promoter extended CGIs present at Polycomb target genes may provide key contact sites for MTF2 **(Figure 4F,G).** Together with the observation that the DNA-binding activity of MTF2 was required for maintenance (**Figure 4C**), these results suggest that MTF2 and its preferential association with extended CGIs at developmental genes are required for inheritance of Polycomb silencing.

### Roles of PRC2 and its read-write capability

We used the heritable silencing system at the *WT1* locus to investigate the hypothesis that PRC2 read-write is required for epigenetic inheritance of Polycomb silencing^32^. We first deleted EED in *5xtetO-H2B-CITRINE* reporter cell lines. In *EED^-/-^* knockout cells, silencing was efficiently established (in +Dox medium), but maintenance of silencing was abolished 8 days after Dox removal (**Figure 5A**). Similarly, deletion of the SUZ12 subunit, a scaffold protein that is required for PRC2 integrity^59^ abolished H3K27me3 and the maintenance of silencing at the *CITRINE* reporter (**Figure S9A, B**). We then attempted to rescue the maintenance defect of *EED^-/-^* cells with either wild-type *HA-EED* or an aromatic cage mutant (F97A, Y148A, Y365A; referred to as *HA-EED-3A*), which does not bind to H3K27me3 and thus cannot allosterically activate EZH2^27,32^ (**Figure 5B, C, Figure S9C**). As reported previously^25^, H3K27me3 was lost in *EED^-/-^* cells (**Figure 5C**). The loss of H3K27me3 in *EED^-/-^* cells was rescued in cells transfected with wild-type *HA-EED* but not the mutant *HA-EED-3A* (**Figure 5C**). The expression of wild-type HA-EED fully rescued the maintenance defect of the *CITRINE* reporter in the *EED^-/-^* cells (**Figure 5D**). Surprisingly, expression of the HA-EED-3A mutant showed a partial rescue in a substantial fraction of cells (∼23%) 8 days after the release of rTetR-CBX7 (**Figure 5D**). To test the possibility that low levels of H3K27me3, which may not have been detected in a whole cell lysate western blot contributed to the partial maintenance rescue observed in HA-EED-3A cells, we performed ChIP-qPCR and ChIP-Seq for H3K27me3. We observed that H3K27me3 was restored at the *WT1* locus during establishment and maintenance in wild-type *HA-EED* cells but was not detectable in *HA-EED-3A* or *EED^-/-^*cells (**Figure 5E, Figure S9D**). Similarly, at the endogenous *PCDH10* Polycomb target gene, H3K27me3 enrichment was restored in *HA-EED* cells but not *HA-EED-3A* cells (**Figure S9D**). Maintenance of silencing in *HA-EED-3A*, but not *EED^-/-^*cells, suggests that the mutant protein performs a maintenance function and should be recruited to the reporter locus. Consistent with this expectation, ChIP-qPCR showed that the HA-EED-3A was recruited to the reporter at the *WT1* locus and remained bound to the locus at low levels during maintenance 4 and 8 days after Dox removal (**Figure S9E, F**). These results indicate that while H3K27me3 and the read-write capability of PRC2 contribute to the stable maintenance of Polycomb-mediated silencing, substantial epigenetic maintenance can occur in the absence of H3K27me3 recognition by the EED subunit of PRC2.

**Figure 5.**
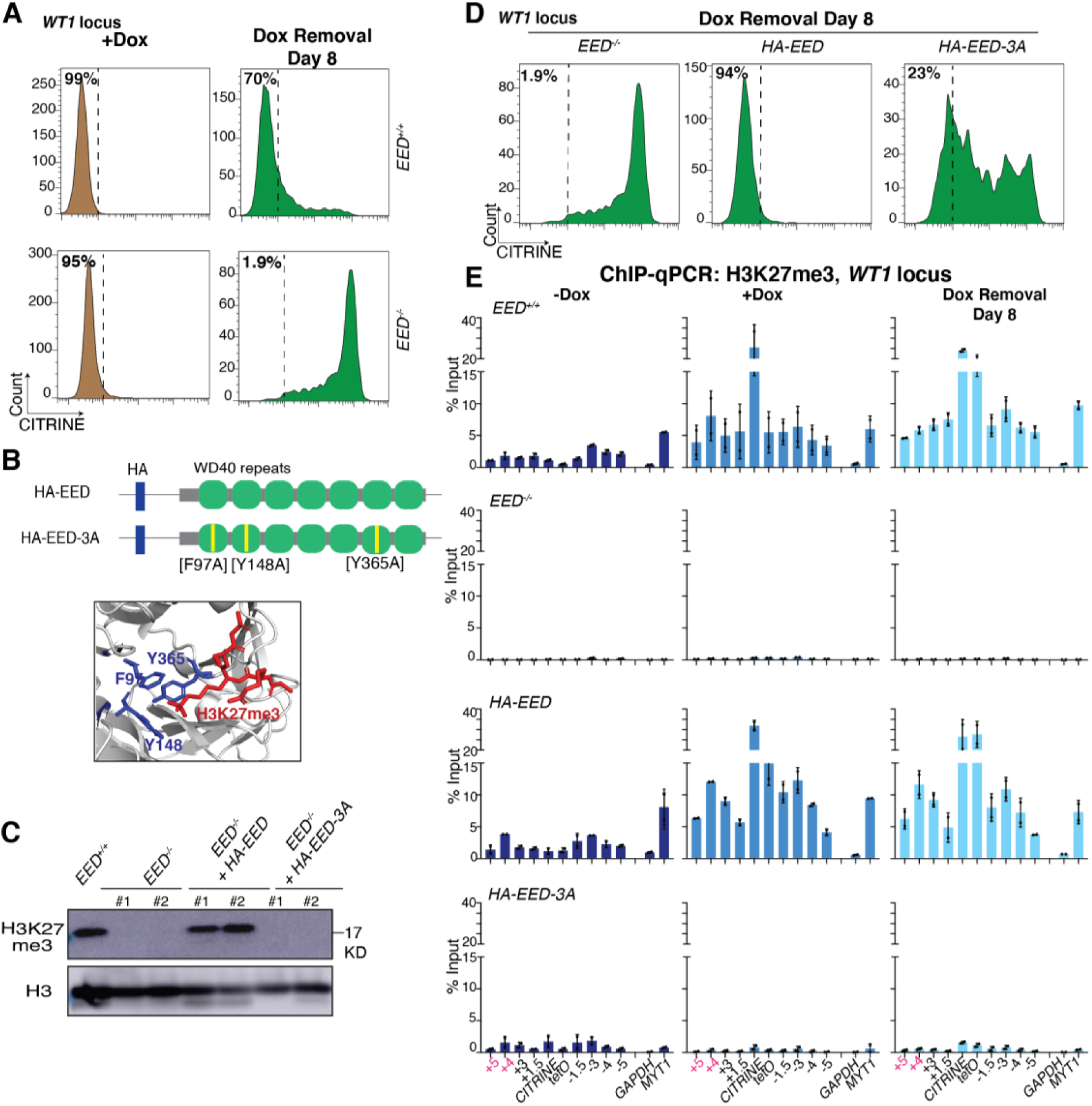
H3K27me3 recognition by PRC2-EED is partly dispensable for epigenetic inheritance of Polycomb silencing. **A**, Flow cytometry histograms showing CITRINE expression, with the reporter inserted at the Polycomb target locus*, WT1,* after establishment of silencing (+Dox) and 8 days after removal of doxycycline (Dox removal) in control (*EED^+/+^*) and *EED* deleted cell line (*EED^-/-^)*. **B**, Schematic diagram showing the wild-type EED (*HA-EED*) and the three residues mutated [F97A,Y148A,Y365A] in the aromatic cage of EED (*HA-EED-3A*) (top). A snapshot of the structure of *EED* aromatic cage with bound H3K27me3 peptide. The *EED* is colored in grey, the F97, Y148 and Y365 residues are in blue, and the H3K27me3 peptide is red (bottom)(PBD3IIW)^27^. **C**, Western blot showing H3K27me3 levels in *EED^+/+^*, *EED^-/-^* and *EED^-/-^* overexpressing *HA-EED* or *HA-EED-3A*. Histone H3 is used as a loading control. **D**, Flow cytometry histograms showing CITRINE expression 8 days after removal of doxycycline (Dox removal) in *EED^-/-^* and *HA-EED* and *HA-EED-3A* cells. **E,** ChIP-qPCR analysis of H3K27me3 enrichment at the reporter locus and surrounding regions before establishment (-Dox), after establishment (+Dox), and 8 days after removal of doxycycline (Dox removal) in *EED^+/+^*, *EED^-/-^*, *HA-EED,* and *HA-EED-3A* cells*. GAPDH* served as a negative control and the native Polycomb-repressed *MYT1* gene as a positive control. Error bars represent standard deviations.

### Key roles for H2AK119ub1 and vPRC1

We next tested the possibility that H3K27me3-independent inheritance of Polycomb silencing is mediated by vPRC1 read-write activity. Consistent with previous findings, deletion of *RING1A* and *RING1B* (*RING1A/B^-/-^)* abolished H2AK119ub1 in our *WT1* reporter cell line (**Figure S10A, B**). The establishment of *CITRINE* reporter silencing was unaffected in *RING1A/B^-/-^* cells, but the maintenance of silencing was greatly diminished (**Figure 6A**). Silencing persisted in a small fraction (16%) of *RING1A/B^-/-^* cells, suggesting that some inheritance could occur independently of H2AK119ub1 (**Figure 6A**). ChIP-qPCR and ChIP-Seq of H3K27me3 showed that during establishment, H3K27me3 was deposited but was greatly decreased by 8 days after release of the rTetR-CBX7 concomitant with de-repression of the *CITRINE* reporter (**Figure 6B, Figure S10C**). These results indicate that in the absence of H2AK119ub1, H3K27me3 is robustly established at the reporter locus, and is likely to recruit downstream factors that silence the reporter gene, but it cannot maintain the silent state in most cells.

**Figure 6.**
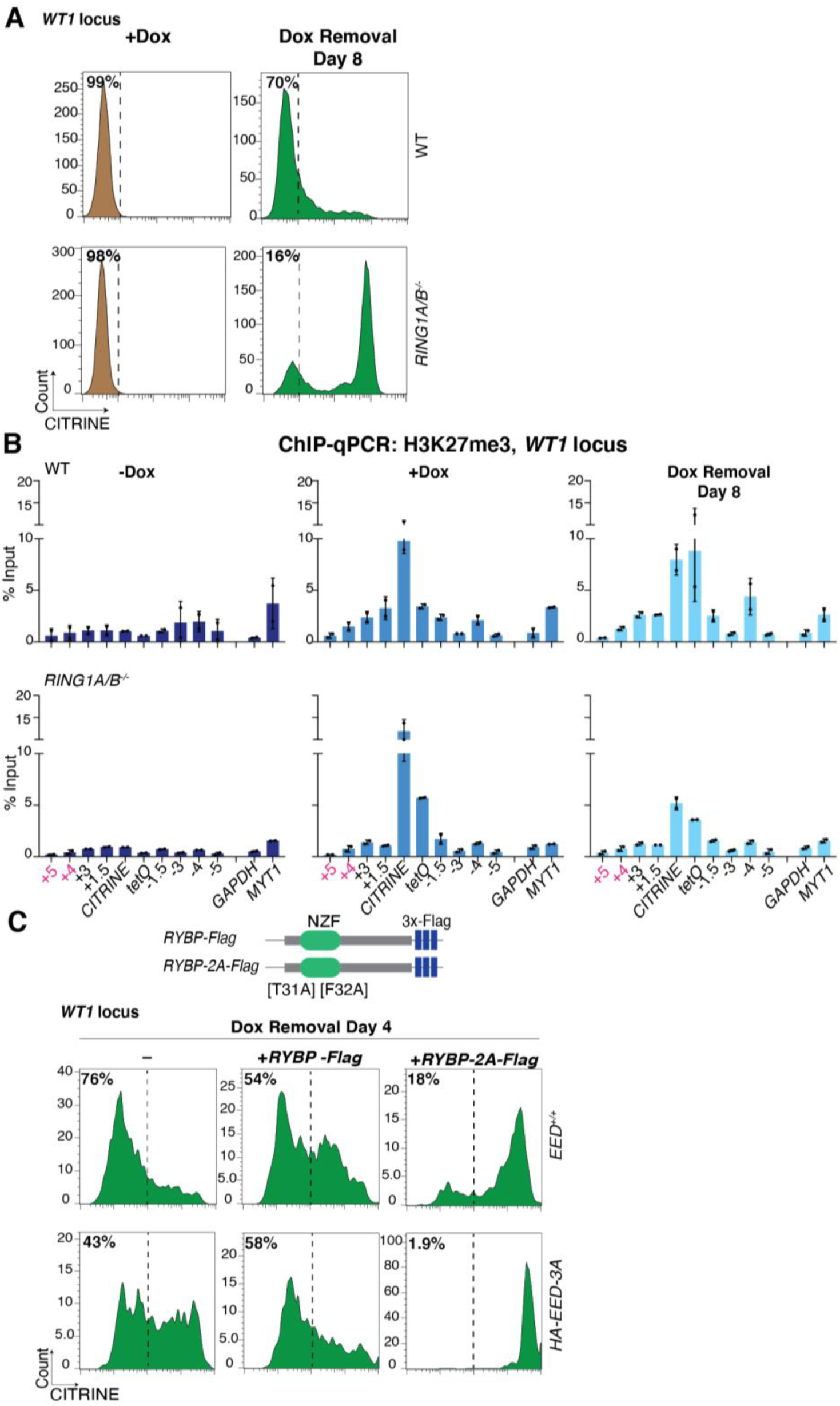
vPRC1 read-write and H2AK119ub1 contribute to epigenetic inheritance of Polycomb silencing independently of H3K27me3 recognition by PRC2. **A,** Flow cytometry histograms showing CITRINE expression after establishment of silencing (+Dox) and 8 days after removal of doxycycline (Dox removal) in wild-type (WT) and *RING1A/B* deleted cell line (*RING1A/B^-/-^)* with the reporter inserted at the Polycomb target locus*, WT1*. **B**, ChIP-qPCR analysis of H3K27me3 enrichment at the reporter locus and surrounding regions before establishment (-Dox), after establishment (+Dox), and 8 days after removal of doxycycline (Dox removal) in Control and *RING1A/B^-/-^*cells. *GAPDH* served as a negative control and the native Polycomb-repressed *MYT1* gene as a positive control. Error bars represent standard deviations. **C**, Schematic diagram showing the wild-type RYBP (*RYBP-Flag*) and the two residues mutated [T31A, F32A] generated in the ubiquitin binding domain of RYBP (*RYBP-2A-Flag*) (top). Flow cytometry histograms showing CITRINE expression 4 days after removal of doxycycline (Dox removal) in *EED^+/+^*and *HA-EED-3A* cells overexpressing RYBP-Flag or RYBP-2A-Flag. (-) refers to non-transfected cells. (bottom).

We next tested the hypothesis that the PRC2 complex containing the mutant HA-EED-3A promotes maintenance of silencing by helping propagate PRC1-mediated H2AK119ub1. In support of this hypothesis, during both the establishment (+Dox) and maintenance (Dox removal) phases, H2AK119ub1 was enriched at the reporter locus in cells with the *HA-EED-3A* construct to similar levels in in *EED^+/+^* cells (**Figure S10D**). H2AK119ub1 is recognized by the RYBP/YAF2 subunit of vPRC1, which allosterically activates RING1A/B ubiquitination activity^36,37^. To test whether this H2AK119ub1 read-write contributes to the inheritance of silencing, we overexpressed either wild-type RYBP-Flag or a ubiquitin binding mutant RYBP (T31A,F32A, called RYBP-2A-Flag)^36,38^ in *EED^+/+^* and *HA-EED-3A* cells with the *CITRINE* reporter at *WT1* (**Figure 6C, Figure S10E**). We reasoned that the mutant RYBP may behave as a dominant negative by competing with wild-type RYBP for incorporation into vPRC1. Overexpression of RYBP-2A-Flag, but not wild-type RYBP-Flag, greatly reduced the maintenance of silencing in *EED^+/+^* cells and abolished the partial maintenance phenotype of *HA-EED-3A* cells (**Figure 6C**). These results demonstrate that vPRC1 read-write of H2AK119ub1 contributes to epigenetic maintenance of Polycomb silencing independently of H3K27me3. They further uncover a role for EED/PRC2 in this inheritance that is independent of H3K27me3 and its recognition.

## Discussion

In this study, we show that newly established domains of Polycomb repression at developmental loci can be epigenetically inherited for many cell divisions (**Figure 7**). This inheritance occurs without any alteration in the trans-acting TFN that regulates cell type-specific gene expression, demonstrating that Polycomb modifications establish a cis memory that cannot be readily erased by the TFN. Importantly, we show that in contrast to developmental loci, newly established Polycomb domains near ubiquitously expressed genes are not heritable, indicating that the inheritance of Polycomb silencing is DNA context-dependent. We further demonstrate that the DNA binding activity of a PRC2 accessory subunit, MTF2, the EED and SUZ12 subunits of PRC2, and the RING1A/B E3 H2AK119 ubiquitin ligase subunits of PRC1 are required for inheritance. However, although deletion of PRC2 subunits abolishes inheritance, indicating a requirement for intact PRC2, disruption of H3K27me3 recognition by the EED subunit of PRC2 does not eliminate inheritance. H3K27me3-independent inheritance requires the RYBP-vPRC1 read-write activity, which may mediate H2AK119ub1 inheritance in the absence of H3K27me3 (**Figure 7**). Below we discuss the implications of these results for mechanism(s) that promote heritable Polycomb silencing.

**Figure 7.**
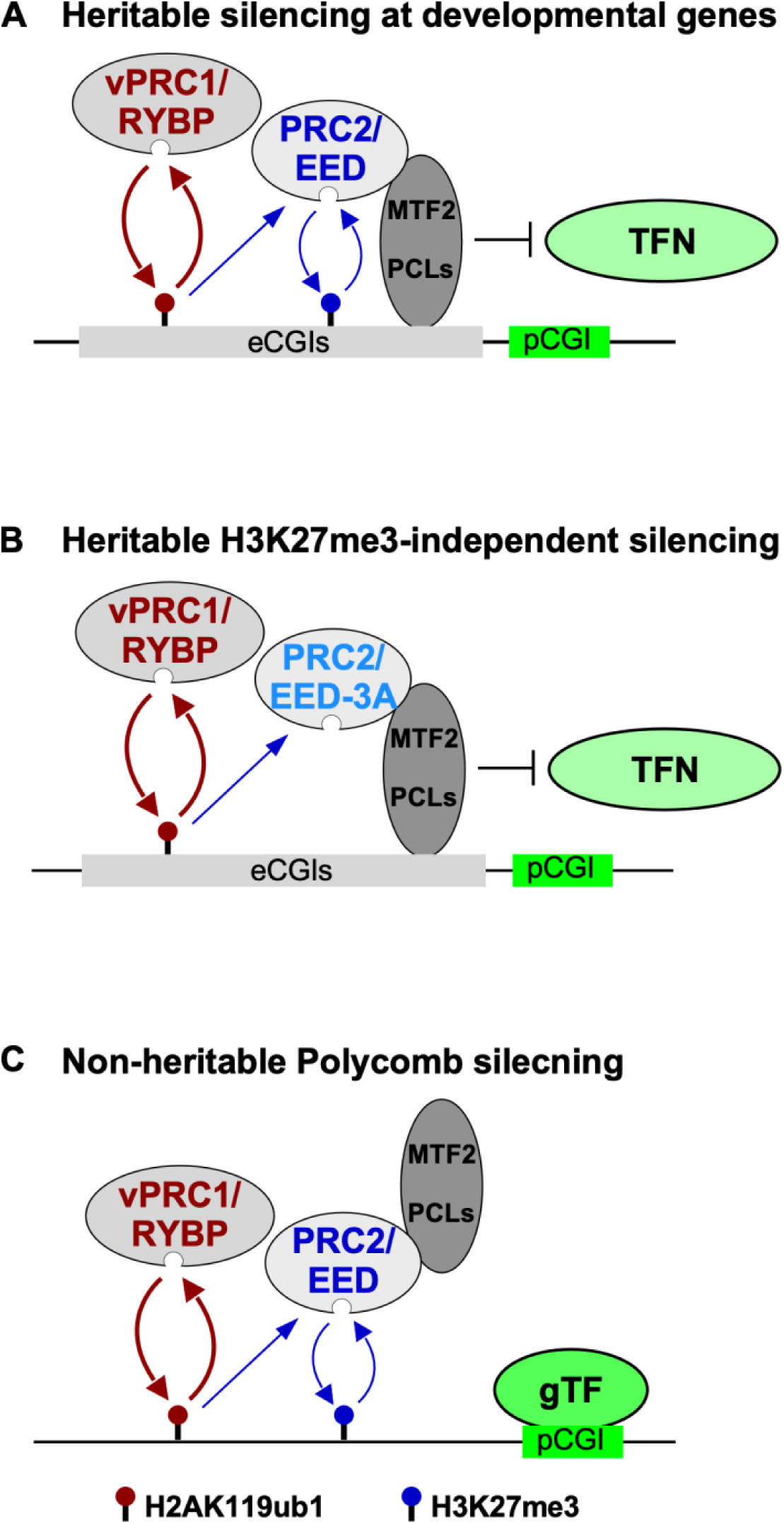
Heritable Polycomb silencing is locus- and H2K119ub1-dependent but only partially H3K27me3-dependent. **A,** At developmental genes, H2AK119ub1 and H3K27me3 positive feedback loops are established by the vPRC1 and PRC2 and work together with MTF2/PCL, which bind to extended CpG islands (eCGIs), to promote heritable silencing and prevent inappropriate TFN-mediated autoactivation. **B,** In EED-3A cells, which lack H3K27me3, Polycomb silencing at a developmental gene can be inherited by an H2AK119ub1- and MTF2-dependent mechanism. **C,** Polycomb silencing induced near ubiquitously expressed housekeeping genes is not heritable due to the absence of eCGIs.

### Restricted Epigenetic Inheritance of Polycomb Silencing

Our findings demonstrate that the inheritance of Polycomb silencing is DNA context-dependent. Once established, Polycomb memory can persist at developmental genes that are regulated by unperturbed cell type-specific transcription factors. However, Polycomb domains established near transcriptionally active ubiquitously expressed genes are unstable, being lost within 2 cell divisions after removal of the rTetR-CBX7 initiator. Thus the inheritance of Polycomb silencing at mammalian genic loci depends on specific sequence features of developmental genes and in addition may be antagonized by constitutively ON enhancers at housekeeping genes. These features are likely to involve the higher abundance of CGIs at developmental genes compared to non-target genes such as ubiquitously expressed genes^48,57,58^. Consistent with this hypothesis, we found that point mutations that disrupt the ability of the PRC2 accessory subunit MTF2 to recognize and bind to CG-rich DNA strongly impaired Polycomb inheritance. The importance of MTF2-mediated CGI recognition in inheritance of Polycomb silencing at developmental genes is additionally supported by our observation that, after the establishment of silencing, both MTF2 and H3K27me3 were enriched at the more extended CGIs of developmental genes but not the single promoter-associated CGI of the ubiquitously expressed *TFRC* gene. Specific DNA sequences have also been shown to be required for inheritance of silent chromatin in *Drosophila* and the fission yeast *S. pombe*^60–63^. In *Drosophila*, PREs are composite DNA elements that are required for both the establishment and inheritance of H3K27me3 domains and silencing of the *HOX* gene clusters^64–66^. Similarly, in *S. pombe*, composite DNA elements called maintainers are essential for maintenance of H3K9me3 domains unless an anti-silencing factor that promotes H3K9me3 demethylation is deleted^62,63,67^. Our findings challenge DNA sequence-independent models of Polycomb inheritance based solely on the read-write positive feedback associated with Polycomb complexes and instead suggest that specific DNA sequences, maintainers in *S. pombe*, PREs in *Drosophila*, and extended CGIs in mammals, act as cis memory modules that serve direct and broadly conserved roles in epigenetic inheritance of different types of silent chromatin. Since both the H3K27me3- and CGI-binding activities of PRC2 are required for inheritance of a newly established silencing, CGIs are likely to promote inheritance by participating in cooperative recruitment of PRC2 (**Figure 7**). CGIs may similarly participate in cooperative recruitment of vPRC1, which can bind to H2AK119ub1 via its RYBP subunit and has been shown to interact with a CGI-binding protein KDM2B^36–38^.

### H2AK119ub1-mediated Polycomb inheritance

Our results suggest that read-write positive feedback loops associated with both the PRC1 and PRC2 complexes are required for robust inheritance of Polycomb silencing (**Figure 7**). However, while intact PRC2 is required for the inheritance of silencing, the ability of its EED subunit of PRC2 to recognize H3K27me3 and promote H3K27 methylation is partially dispensable. This is surprising since current models of Polycomb inheritance are based on this well-documented write and read mechanism. In the absence of H3K27me3, we found that PRC1-mediated H2AK119 ubiquitination and the ability of the RYBP subunit of vPRC1 to recognize H2AK119ub1 are absolutely essential for heritable silencing. In this regard, the ubiquitin binding domain of RYBP was previously shown to be required for efficient maintenance of H2AK119ub1 at an inducible reporter gene, but its role in epigenetic inheritance remained unclear since genomic integration of the reporter in mESC cells was coupled to histone H3K9 and DNA methylation, which are generally absent at mammalian Polycomb target genes^36^. H2AK119ub1-dependent and H3K27me3-independent inheritance of silencing raise questions about the mechanism of chromatin inheritance during DNA replication. In the current models for inheritance of parental histones during DNA replication, parental histone H3/H4 tetramers are thought to be transferred to newly synthesized daughter DNA strands. Histone H2A/H2B dimers, on the other hand, are thought to be rapidly exchanged and not inherited^68–70^. Our findings raise the possibility that specialized mechanism have evolved to ensure H2AK119ub1/H2B dimer inheritance within Polycomb domains.

## Materials and Methods

### Cell Culture

HEK293FT (ThermoFisher R70007) cells were maintained in DMEM medium (Invitrogen) plus 10% FBS (Invitrogen), 1 mM Glutamine, and 100 µg/ml penicillin-streptomycin following standard culture conditions. To induce the binding rTetR-CBX7 to the *5xtetO* Site, 1 µg/ml doxycycline (Sigma, D9891) was added to the culture medium. The reagents used in this study are listed in Table S1. The cell lines generated in this study are listed in Tables S2-S3.

### Plasmid Construction

Donor Plasmids for insertion of *5xtetO-pEF1-H2B-CITRINE* reporter into the genome were constructed by subcloning *5xtetO-pEF1-H2B-CITRINE-PolyA* from PhiC31-Neo-ins-5xtetO-pEF-H2B-Citrine-ins (Addgene# 78099)^14^ with right and left homology arms (500 bps each) in CloneSmart HCKan Blunt (Gift from Jichuan Zhang [Initiative for Genome Editing and Neurodegeneration core in the Department of Cell Biology at Harvard Medical School], Lucigen# 40704-2). Plasmid with mCherry-2A-rTetR-CBX7 was created by subcloning mCherry-2A-rTetR (Addgene # 78101)^14^ and CBX7 into lentiviral expression vector backbone pLVU-tTR-KRAB (Addgene# 11645)^71^. The CBX7 open reading frame was amplified from pCMV-SPORT6-CBX7 (DFCI Plasmid# HsCD00339744). Rescue Plasmids were constructed by cloning HA or 3X Flag tags fused to EED, MTF2, and RYBP wild-type and mutant cDNAs using Gibson assembly into pdCas9-DNMT3A-2A-PuroR (Addgene# 71667)^72^, replacing pdCas9-DNMT3A. In the case of RYBP, PuroR was replaced with hygromycin. The point mutations in EED, MTF2, and RYBP were generated using IDT gBlocks.

### CRISPR genome editing

sgRNAs for reporter cell line construction, gene knockouts and sequence deletions were designed using the CRISPR design tool in https://benchling.com and/or https://chopchop.cbu.uib.no/ (Table S2). sgRNAs were either *in vitro* transcribed using GeneArt™ Precision gRNA Synthesis Kit (ThermoFisher A29377) and electroporated with Neon Transfection System (ThermoFisher MPK1025), along with donor plasmid and Cas9 protein (Gift from Jichuan Zhang, Initiative for Genome Editing and Neurodegeneration core in the Department of Cell Biology at Harvard Medical School) or cloned into pSpCas9(BB)-2A-Puro (PX459) V2.0 (Addgene plasmid # 62988) and transfected into HEK293FT using Lipofectamine 2000 (ThermoFisher). sgRNAs for rTetR-CBX7 deletion were cloned into LentiCRISPRv2 (Addgene 52961). CITRINE positive cells were sorted into single cell colonies in 96 well plates, genotyped by PCR (genotyping primer sequences are presented in Table S4) and confirmed by Sanger Sequencing (Quintara Bio) or MiSeq (Illumina). Southern Blot using a CITRINE probe was carried out to verify single integration for the reporter cell lines.

### Integration of Rescue Constructs

Rescue Plasmids were transfected into relevant cell lines using Lipofectamine 2000 (Thermofisher). Cells with insertions were selected using Puromycin (ThermoFisher) at 0.6 µg/ml or hygromycin (ThermoFisher) at 200 µg/ml for 2 weeks. See Table S3.

### Western Blot

Whole cell extract was obtained by lysis in RIPA buffer (final concentrations: 150 mM NaCl, 1% triton, 0.5% sodium deoxy-cholate, 0.1% SDS, 50 mM Tris pH 8.0) and histones were extracted using 0.2 N HCl. The protein concentration was determined by the Bradford assay (Biorad). 10-20 µg/lane total protein was electrophoresed on 4–15% Mini-PROTEAN® TGX™ Precast Protein Gels (BioRad) with SDS running buffer and transferred to Polyvinylidene difluoride (PVDF) membranes. The membranes were blocked (5% non-fat dry milk in 1× PBS, 0.1% Tween 20) for 2 hrs and then incubated in 5% non-fat dry milk in 1× PBS (137 mM NaCl, 2.7 mM KCl, 8 mM Na2HPO4, and 2 mM KH2PO4.), 0.1% Tween 20 with the primary antibodies as listed in Table S5 for 2 hours at room temperature or overnight at 4°C. Finally, membranes were incubated with corresponding secondary Licor IRDye antibody (5% non-fat dry milk in 1× PBS, 0.1% Tween 20) and imaged by Odyssey Clx (Licor) or HRP-conjugated secondary antibodies and imaged on autoradiography film/Amersham Imager (GE).

### RT-qPCR

Total RNA was extracted using the RNeasy Mini kit (74104, Qiagen) and reverse transcribed into cDNA using random hexamers (Invitrogen) and reverse transcription kit (18090010, ThermoFisher). cDNA was analyzed using PCR on a QuantStudio 7 Flex Real Time PCR System (Applied Biosystem). PCR parameters were 95°C for 2 min and 40 cycles of 95°C for 15s, 60°C for 30s, and 72°C for 15 s, followed by 72°C for 1 min. All the qPCR data presented were at least two biological replicates and plotted with Prism GraphPad Software with error bars representing standard deviation. Primer sequences are presented in Table S6.

### LentiViral Production and Infection

Plasmids were purified using a MaxiPrep DNA isolation Kit (Qiagen). For virus packaging, we used psPAX2 (Addgene# 12260) and pMD2.G (Addgene# 12259) which were transfected into HEK293FT cells using Lipofectamine 2000 (Invitrogen). Medium containing the viral particles was collected 72 hr after transfection and viral particles were concentrated using the PEG-it Virus precipitation solution (SBI LV810A-1). Cells were transduced with the virus for 48 h in the presence of 4 µg/ml polybrene (Sigma H9268).

### Fluorescence Imaging

Cells were plated on chamber slides (ThermoFisher 154526PK). Cells were first washed with PBS, fixed with 4% paraformaldehyde in PBS for 5 mins and permeabilized with PBS/0.25% Triton X-100 at room temperature for 5 mins. Cells were mounted with VECTASHIELD HardSet Antifade Mounting Medium with DAPI (Vector Labs) and imaged in the DAPI, YFP and RFP channels using a widefield microscope (Nikon Ti2) equipped with a 40× objective lens (Nikon Imaging Center at Harvard Medical School). Images were post-processed with Image J^73^.

### Fluorescence-Activated Cell Sorting and Analysis

Cells were made into single cell suspension using 0.05% Trypsin (Invitrogen) and suspended in HEK293FT culture medium. Samples for analysis were collected with LSR Fortessa (BD Biosciences, Dana Farber Flow Cytometry core) or FACs Calibur (BD Biosciences, Department of Cell Biology at Harvard Medical School). The GFP channel was used for CITRINE detection. Samples were sorted with M AriaII (BD Biosciences, Immunology Flow Cytometry core at Harvard Medical School). Data was analyzed with FlowJo™ Version 10.5.3 (Ashland, OR: Becton, Dickinson and Company; 2021). Experiments were performed with at least two biological replicates.

### ChIP-qPCR and ChIP-Seq

ChIP was performed as previously described with minor modifications ^74^. Cells for ChIP were cultured in 15 cm plates (∼10 million cells). Cell pellets were first washed with cold PBS, crosslinked at room temperature with 1% formaldehyde (ThermoFisher Scientific) for 8 min. Crosslinking reactions were quenched by addition of 125 mM glycine for 10 min. Cell were then resuspended in Swelling Buffer (25 mM Hepes pH 7.8, 1.5 mM MgCl_2_, 10 mM KCl, 0.1% NP-40, 1 mM DTT) followed by Dounce homogenization. Nuclei were pelleted by centrifugation and then resuspended in sonication buffer (0.1% SDS, 1 mM EDTA and 10 mM Tri-HCl pH 8.0). The nuclei were sonicated to shear chromatin into ∼200-500 bp fragments using a Covaris E220. Sonicated samples were diluted with ChIP dilution buffer (0.1% SDS, 1 mM EDTA and 10 mM Tri-HCl pH 8.0, 1% Triton X-100, 150 mM NaCl). Diluted samples were centrifuged at 13,000 rpm for 10 min. The supernatant was used for immunoprecipitation using antibodies and 25 μl protein A/G beads for 12-16 h at 4°C (see Table S5 for antibodies). For H3K27me3 ChIP-Seq, *Drosophila* S2 chromatin (Active Motif# 53083) and histone H2Av antibody (Active Motif# 61686) were added as spike-in controls. ChIP-Seq samples for Flag antibody do not have spike-in controls. The beads were washed twice with high salt wash buffer A (50 mM Hepes pH 7.9, 500 mM NaCl, 1 mM EDTA, 1% Triton, 0.1% Sodium deoxycholate, and 0.1% SDS), twice with wash buffer B (20 mM Tris-HCl pH 8.0, 250 mM LiCl, 1 mM EDTA, 0.5% Sodium deoxycholate, 0.5% NP-40) and twice with 1X TE (10 mM Tris-HCl, 1mM EDTA). The bound chromatin fragments were eluted with elution buffer (50 mM Tris pH 8.0, 1 mM EDTA, 50 mM NaHCO_3_,1% SDS) twice for 10 min each at 65°C. Eluted DNA-proteins complexes were incubated overnight at 65°C to reverse crosslinks. RNAase A followed by Proteinase K was then added to digest RNA and protein. DNA was further purified using phenol chloroform/PCR Purification Kit (QIAGEN) and analyzed by PCR on a QuantStudio 7 Flex Real Time PCR System (Applied Biosystem). PCR parameters were 95°Cfor 2 min and 40 cycles of 95°C for 15s, 60°C for 30s, and 72°C for 15 s, followed by 72°C for 1 min. All the ChIP-qPCR data presented include at least two biological replicates. Primer sequences are in Table S7. Results were plotted with Prism GraphPad Software with error bars representing standard deviation.

For ChIP-seq, sequencing libraries were constructed using TruSeq DNA sample Prep Kits (Illumina) and adapter dimers were removed by 2% agarose and Tris-acetate-EDTA gel electrophoresis. Size selected and purified DNA libraries were sequenced on an Illumina NextSeq 500 machine (Bauer core facility at Harvard University) to obtain 75 bp single-end reads. ChIP-seq reads were quality controlled with fastqc (v0.11.5) and mapped to the human genome reference (Custom *5xtetO-H2B-CITRINE* Reporter Inserted at Chr11-hg19 near *WT1* or Custom *5xtetO-H2B-CITRINE* Reporter Inserted at Chr3-hg19 near *TFRC*) and *Drosophila* (dm3) using bowtie2 (v2.2.9) with default parameters. Scale factor was calculated as previously described to normalize H3K27me3 signal ^75^. Bam files were generated with samtools 1.3.1, which was followed by making bigwig files with deeptools (v/3.0.2)^76,77^. Reads were normalized with scale factor for H3K27me3 or RPGC for Flag with deeptools (v/3.0.2) bamCoverage function. ChIP-Seq tracks were visualized in Integrative Genomics Viewer (IGV). Publicly available source data used for this study are listed in Table S8.

### Data Availability

Software and algorithms used in this study are listed in Table S9. The raw and processed high-throughput sequencing data have been deposited at NCBI Gene Expression Omnibus under ID code GEO: GSE209986.

## Acknowledgments

We thank B. Bernstein, W. Feng, S. Parhad, R. Sotomayor, D. Yaghoubi, A. Tatarakis, and J. Yu for comments and members of the Moazed lab for helpful discussions. We thank the Bauer Core Facility at Harvard University for high-throughput sequencing, H. Saini for advice on bioinformatics analysis, Jiuchun Zhang of the Initiative for Genome Editing and Neurodegeneration at the Department of Cell Biology at Harvard Medical School for advice and reagents for CRISPR-Cas9 deletions, the Nikon Imaging Center at the Department of Cell Biology at Harvard Medical School for access to microscopes and advice, and the Immunology Flow Cytometry Core at Harvard Medical School and the Dana Farber Flow Cytometry core for access to Flow Cytometry equipment. D.M. is an investigator of the Howard Hughes Medical institute.

## Author contributions

T.A.S. and D.M. conceived the study and designed experiments. T.A.S. performed all experiments. T.A.S. and D.M. wrote the manuscript.

---

**Figure S1.**
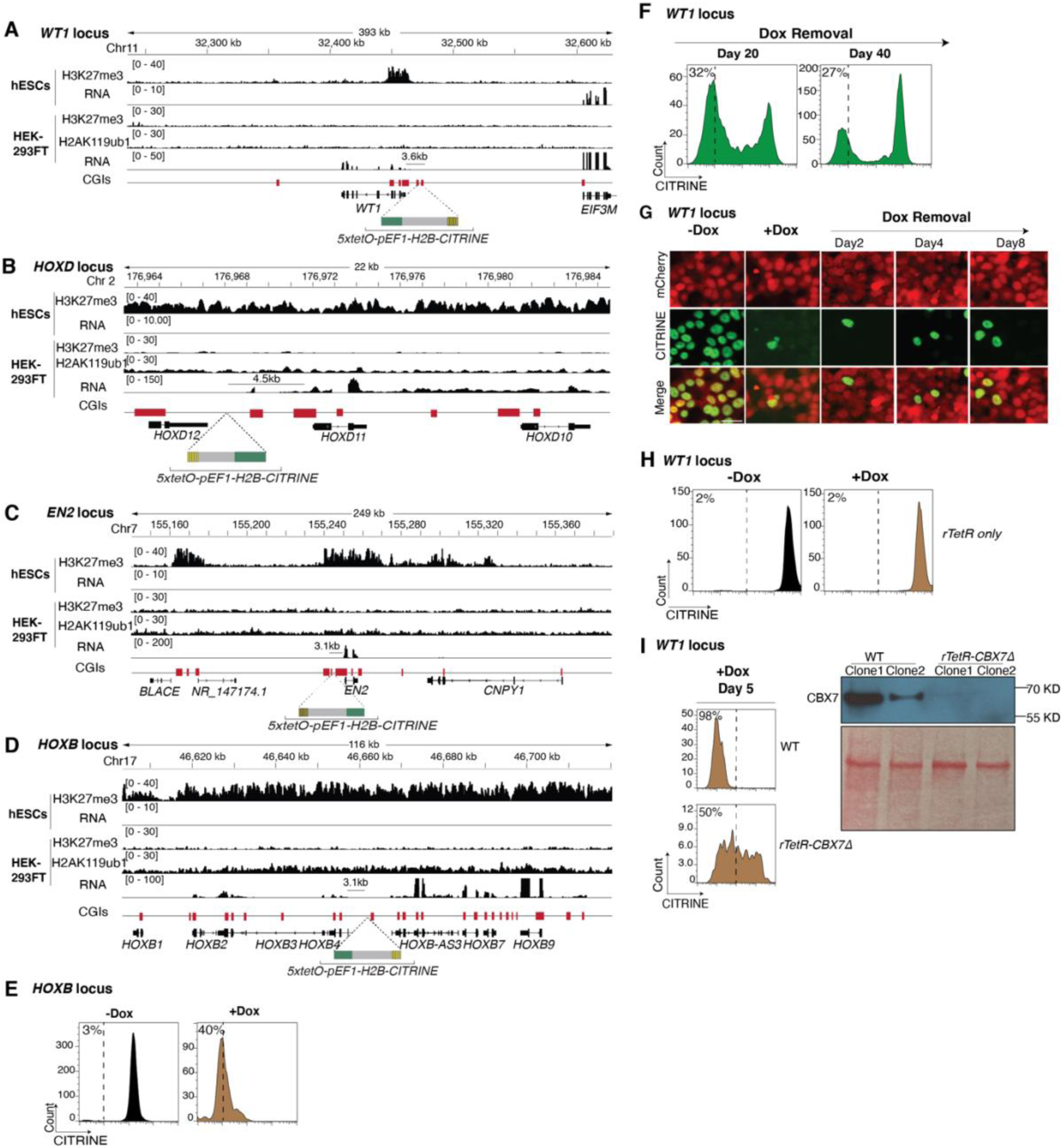
Chromatin features and additional control experiments at reporters inserted near developmental genes. Genome browser tracks showing publicly available chromatin features of the *WT1* (**A**), *HOXD* (**B**), *EN2* (**C**) and *HOXB* (**D**) loci in human embryonic stem cells (hESCs) and HEK293FT cells (see Table S8 for references). Chromosome coordinates (top) and the site of insertion of the *5xtetO-pEF1-H2B-CITRINE* reporter (bottom) are indicated. **E,** Flow cytometry histograms showing CITRINE expression before and after establishment of silencing (-Dox and +Dox) near *HOXB4*. Percentages (%) indicate the fraction of CITRINE negative cells. **F,** Flow cytometry histograms showing CITRINE expression at the *WT1* locus at 20 and 40 days after removal of doxycycline (Dox removal). Percentages (%) indicate the fraction of CITRINE negative cells. **G,** Representative fluorescence images showing CITRINE expression at the *WT1* locus before and after establishment of silencing (-Dox and +Dox) and at the indicated days after removal of doxycycline (Dox removal). mCherry indicates the expression of rTetR-CBX7 in the reporter cell line. **H,** Flow cytometry histogram showing CITRINE expression at the *WT1* locus before and after doxycycline addition (-Dox and +Dox) in a cell line expressing rTetR-only. Percentages (%) indicate the fraction of CITRINE negative cells. **I,** Flow cytometry histograms showing CITRINE expression at the *WT1* locus in cells with rTetR-CBX7 excised after establishment with doxycycline addition (+Dox) and western blot showing the expression levels of rTetR-CBX7 before and after excision.

**Figure S2.**
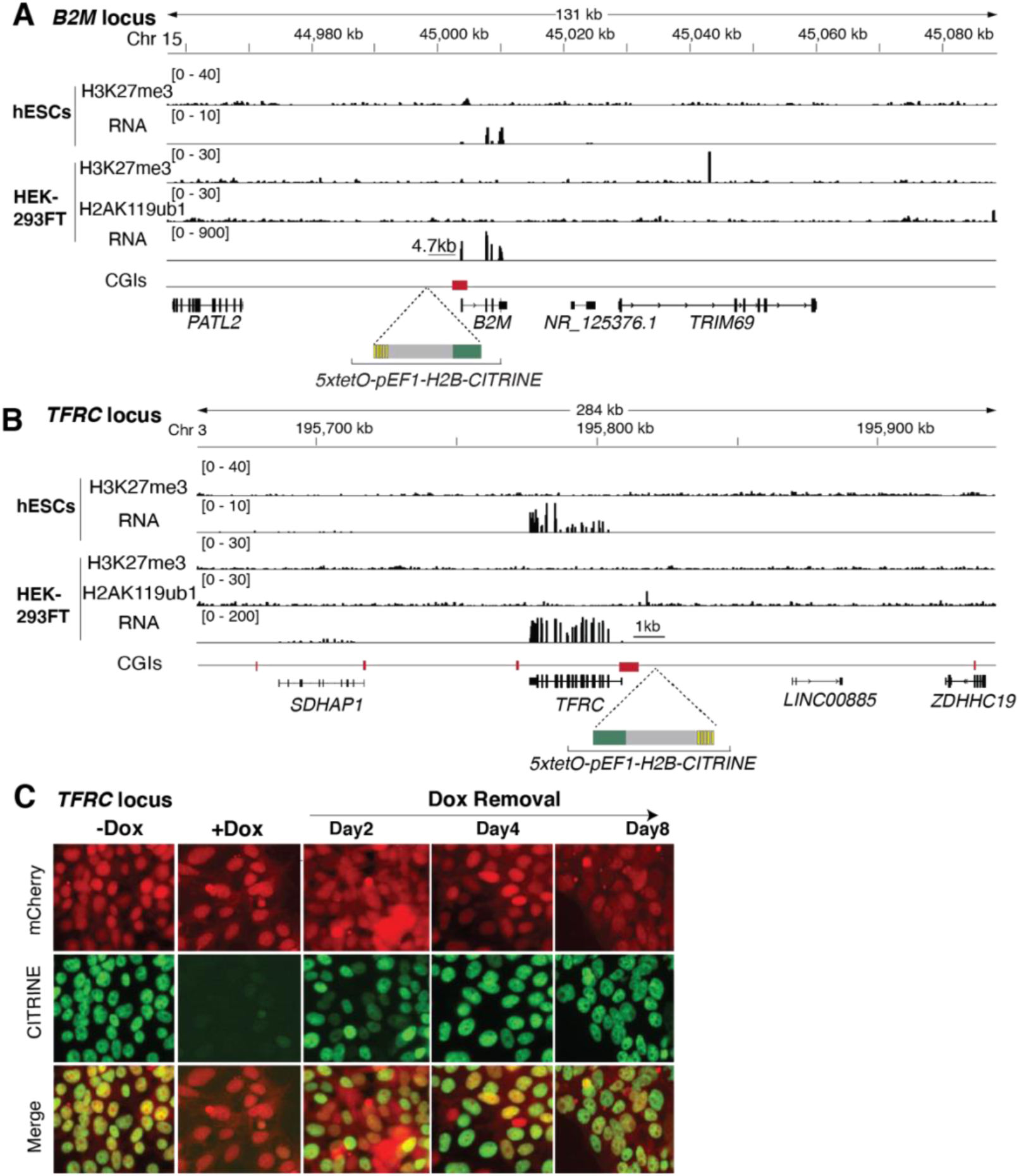
Chromatin features and additional control experiments at reporters inserted near ubiquitously expressed genes. Genome browser tracks showing publicly available chromatin features of the *B2M* (**A**) and TFRC (**B**) loci in human embryonic stem cells (hESCs) and HEK293FT cells (see Table S8 for references). Chromosome coordinates (top) and the site of insertion of the *5xtetO-pEF1-H2B-CITRINE* reporter (bottom) are indicated. **C,** Representative fluorescence images showing CITRINE expression before and after establishment of silencing (-Dox and +Dox) and at the indicated days after removal of doxycycline (Dox removal) at the reporter locus inserted near *TFRC*. mCherry indicates the expression of rTetR-CBX7 in the reporter cell line.

**Figure S3.**
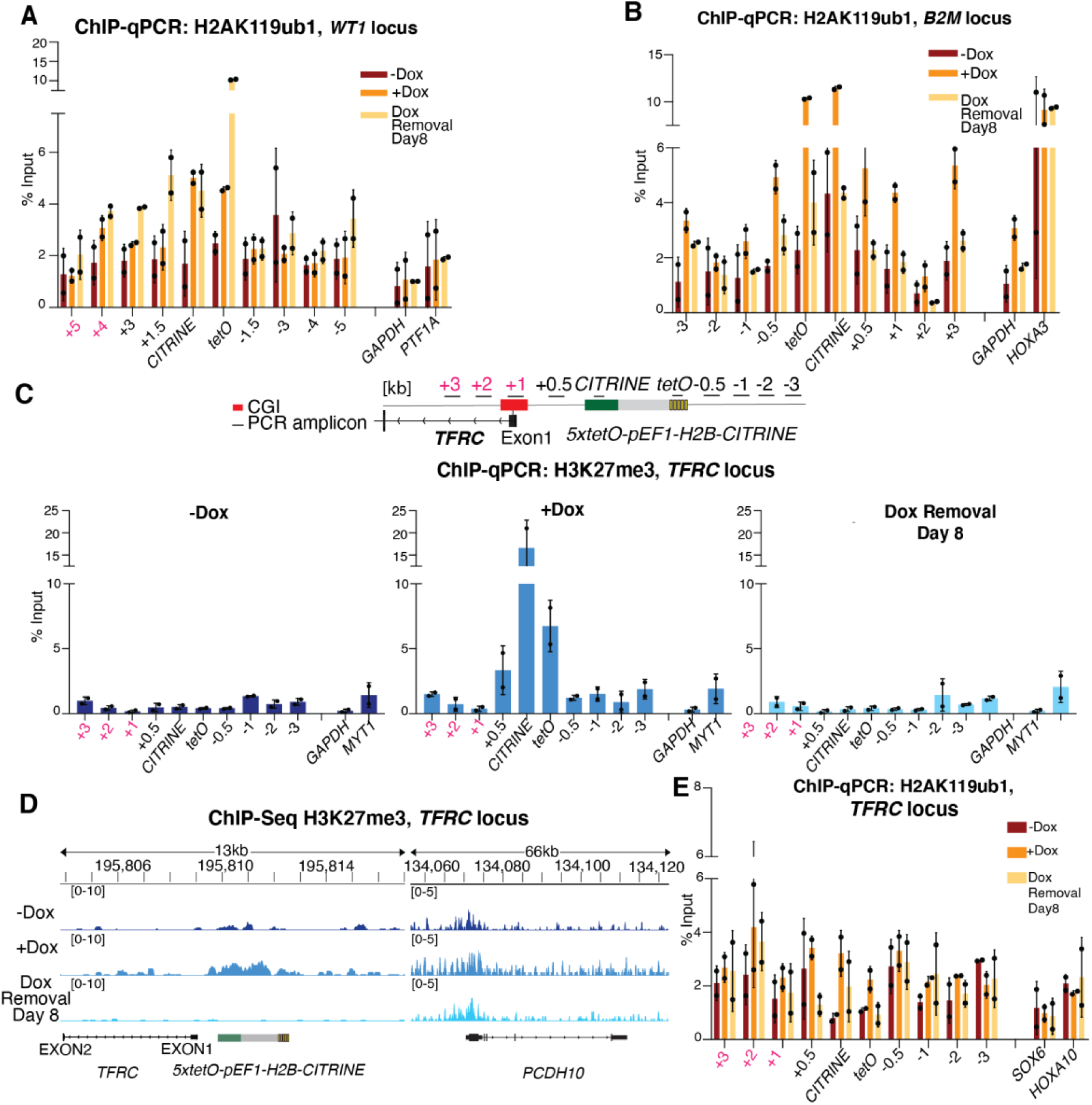
H2AK119ub1, and H3K27me3 are established at *WT1, B2M* and *TFRC* loci, but only inherited at the *WT1* locus. **A,** ChIP-qPCR analysis of H2AK119ub1 enrichment at the reporter locus and surrounding sequences before establishment (-Dox), after establishment (+Dox), and at 8 days after removal of doxycycline (Dox removal) near the *WT1* locus. *GAPDH* served as a negative control and *PTF1A* as a positive control. Error bars represent standard deviations. See Figure 2 for primer maps. **B**, Same as **A** but at the *B2M* locus. See Figure 2 for primer maps. **C,** ChIP-qPCR analysis of H3K27me3 enrichment at the reporter locus and surrounding sequences before establishment (-Dox), after establishment (+Dox), and at 8 days after removal of doxycycline (Dox removal) near the *TFRC* locus. *GAPDH* served as a negative control and *MYT1* as a positive control. Error bars represent standard deviations. **D**, Genome browser snapshots of H3K27me3 ChIP-seq reads at the reporter locus near *TFRC* and the endogenous Polycomb-silenced *PCDH10* gene. **E,** Same as **C**, but showing H2AK119ub1 ChIP-qPCR.

**Figure S4.**
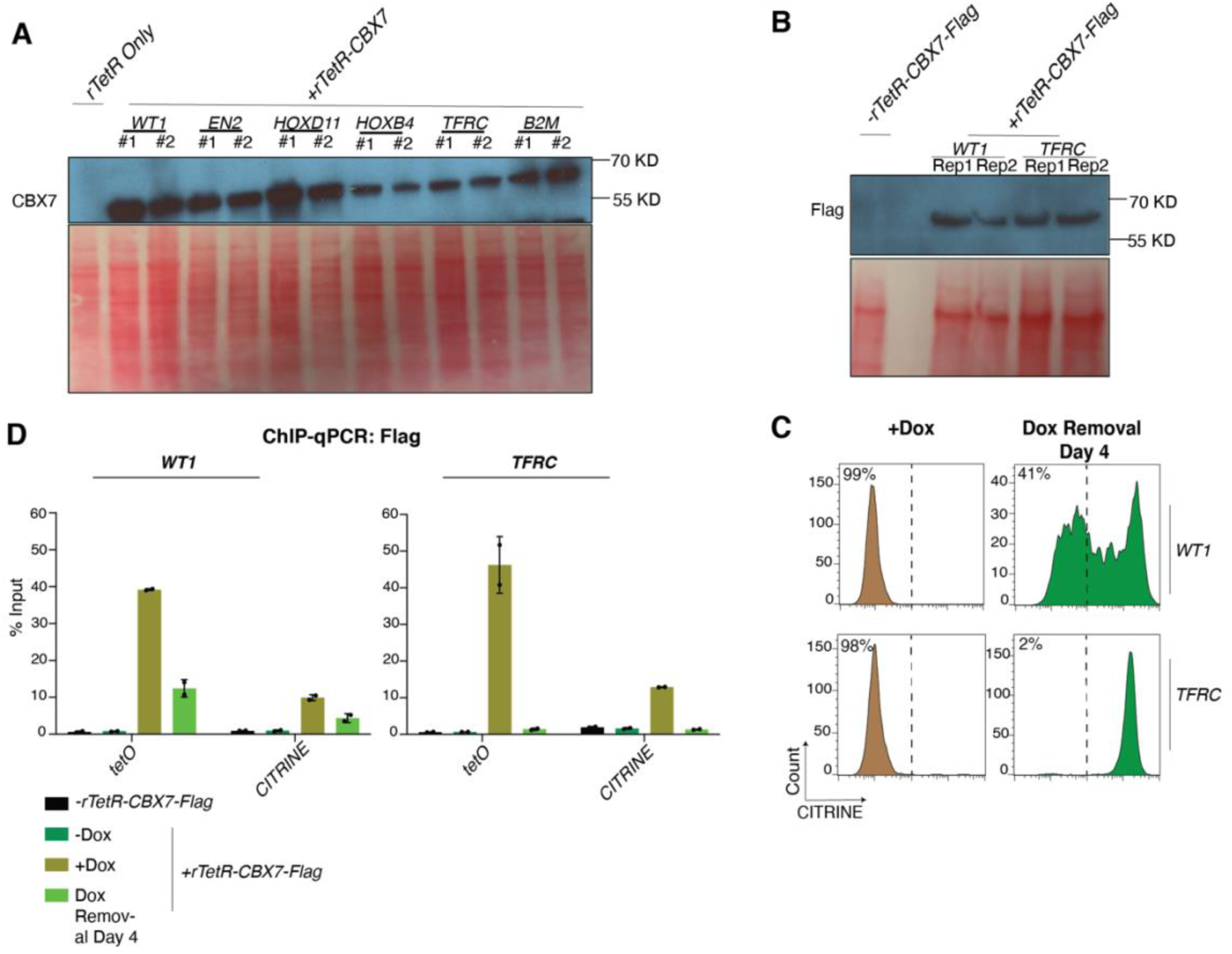
rTetR-CBX7 is expressed and bound to the *tetO* arrays at similar levels at the *WT1* and *TFRC* loci. **A,** Western blot showing the expression levels of rTetR-CBX7 in the reporter cell lines used in this study (two replicates for each cell line are presented). **B,** Western Blot showing the expression levels of rTetR-CBX7-Flag expressed in the *WT1* and *TFRC* reporter cell lines (top). **C**, Flow cytometry histograms showing CITRINE expression after establishment of silencing with doxycycline addition (+Dox) and at 4 days after removal of doxycycline (Dox removal) (bottom) in the cell lines in **B**. Percentages (%) indicate the fraction of CITRINE negative cells. **D,** ChIP-qPCR analysis of Flag at the reporter loci at *WT1* and *TFRC* before establishment (-Dox), after establishment (+Dox), and at 4 days after removal of doxycycline (Dox removal). Reporter cell lines without expression of rTetR-CBX7-Flag served as controls. Error bars represent standard deviations.

**Figure S5.**
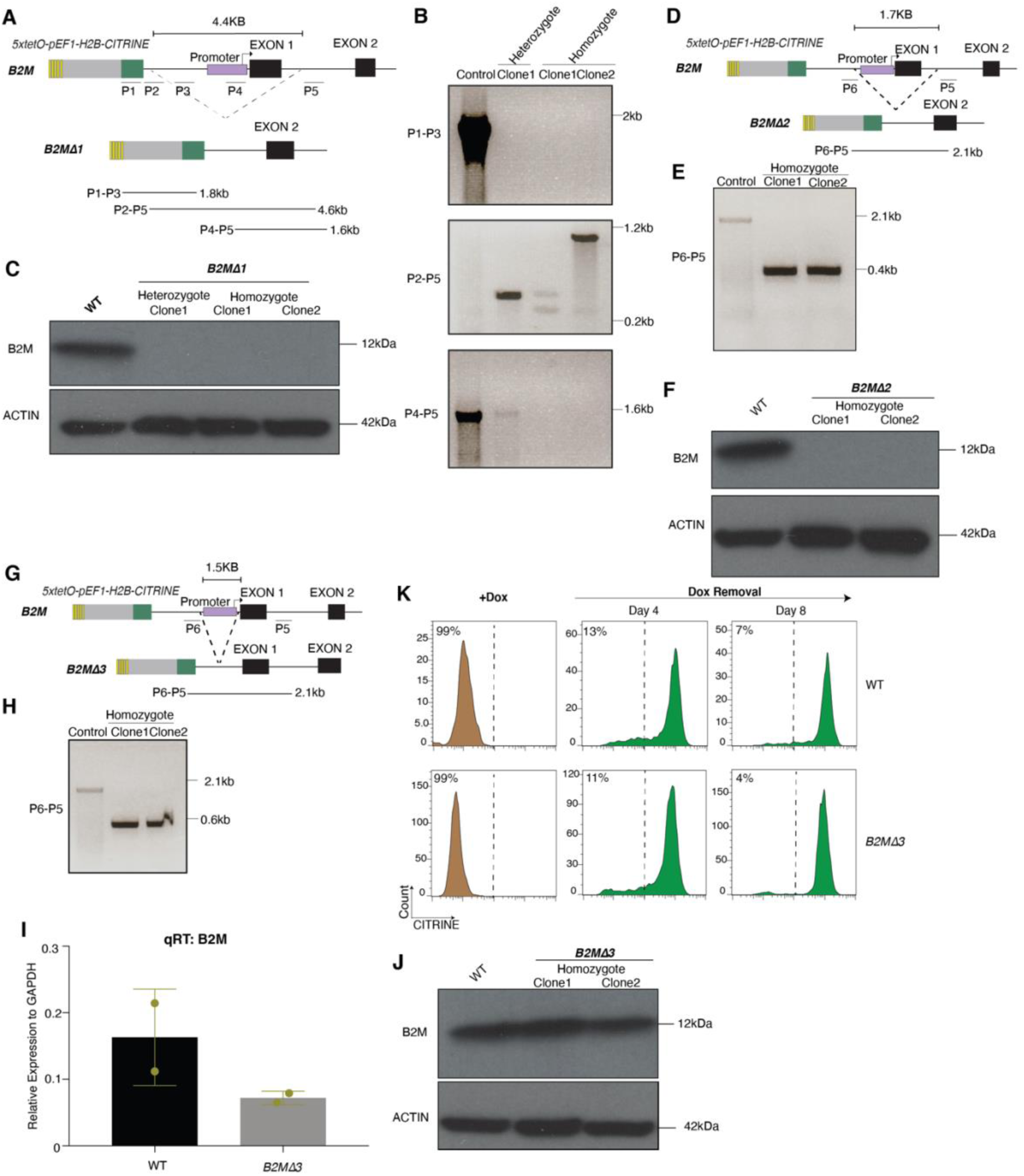
Generation of promoter region deletions at the endogenous *B2M* gene and their effect on epigenetic inheritance of Polycomb silencing. **A,** Schematic diagram of PCR genotyping primers indicating the sequence deletion at *B2MΔ1*. See Figure 3 for genome browser view of the *B2M* locus. **B,** PCR genotyping validating the indicated sequence deletion at *B2MΔ1*. **C,** Western Blot showing the expression B2M in wild-type control (WT) and *B2MΔ1* cell lines. ACTIN serves as a loading control. **D,** same as **A** but showing the location of primers for genotyping *B2MΔ2*. **E**, PCR validation of *B2MΔ2* clones. **F**, Western blots showing B2M protein levels in *B2MΔ2* cell lines. **G**, same as **A** but showing the location of primers for genotyping *B2MΔ3*. **H,** PCR validation of *B2MΔ3* clones. **I**, qRT-PCR showing the steady state B2M mRNA levels in *B2MΔ3* clones, normalized to GAPDH RNA. Error bars represent standard deviation. **J,** Western blots showing B2M protein levels in *B2MΔ3* clones. **K,** Flow cytometry histograms showing CITRINE expression after establishment of silencing (+Dox) and at the indicated days after removal of doxycycline (Dox removal) in WT and *B2MΔ3* cell lines. Percentages (%) indicate the fraction of CITRINE negative cells.

**Figure S6.**
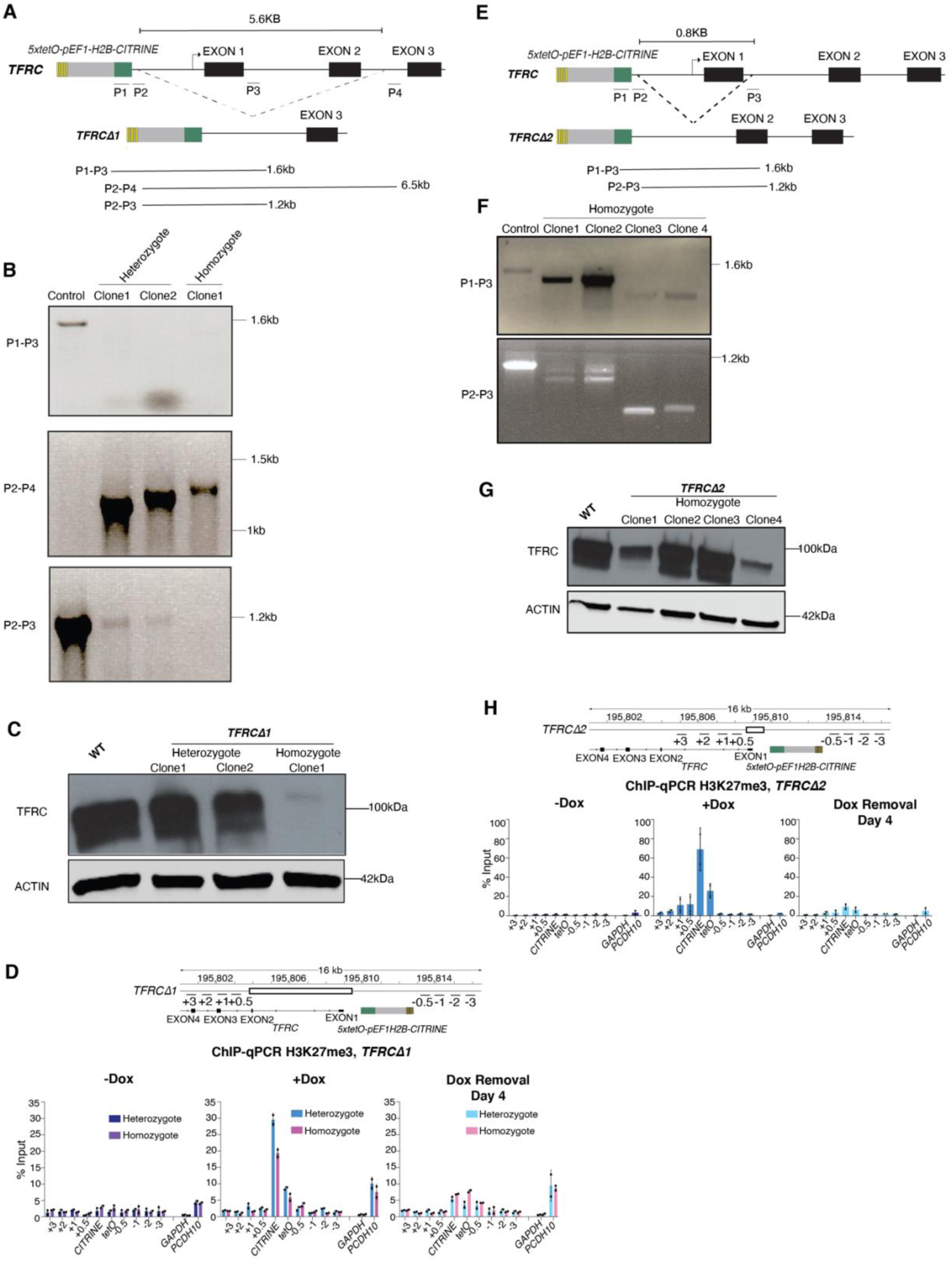
Generation of promoter region deletions at the endogenous *TFRC* gene. **A,** Schematic diagram of PCR genotyping primers used for validating the sequence deletion at *TFRCΔ1* clones. See Figure 3 for genome browser view of the *TFRC* locus **B,** PCR assays for genotyping the indicated sequence deletion at *TFRCΔ1* clones. **C,** Western Blot showing the expression levels of TFRC in wild-type control (WT) and *TFRCΔ1* cell lines. ACTIN serves as a loading control**. D,** ChIP-qPCR analysis of H3K27me3 enrichment at the reporter locus and surrounding regions before establishment (-Dox), after establishment (+Dox), and 4 days after removal of doxycycline (Dox removal) in *TFRCΔ1* heterozygote and homozygote clones. *GAPDH* served as a negative control and the native Polycomb loci *PCDH10* gene as positive control. Error bars represent standard deviation. **E,** Schematic diagram of PCR genotyping primers used for validating the sequence deletion at *TFRCΔ2* clones. **F,** PCR assays for genotyping the indicated sequence deletion at *TFRCΔ2* clones. **G,** Western Blot showing the expression levels of TFRC in wild-type control (WT) and *TFRCΔ2* cell lines. **H,** same as **D** but for *TFRCΔ2* cells.

**Figure S7.**
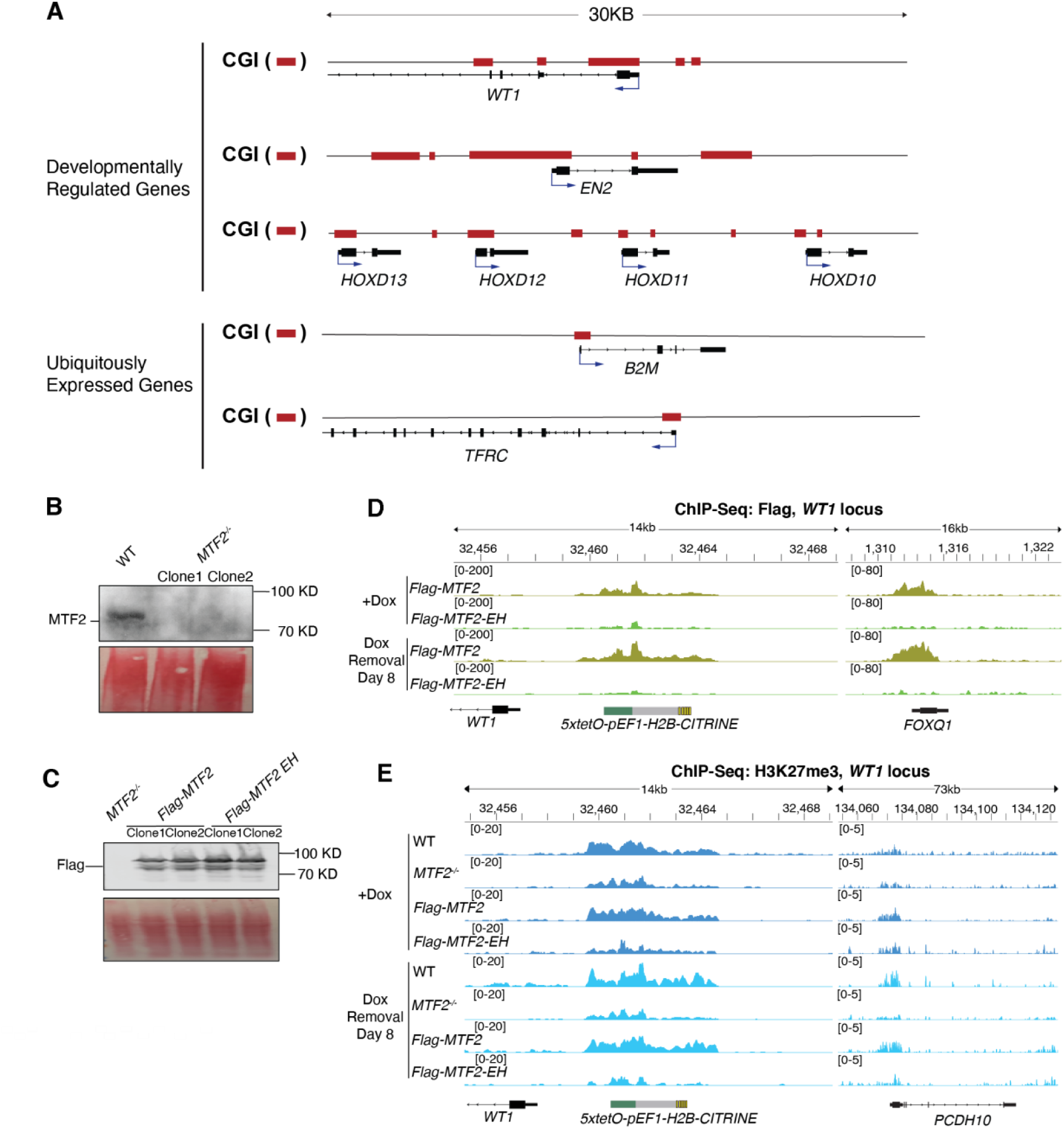
Mutations in Extended Homology Domain of MTF2 lead to loss of its chromatin localization and loss of H3K27me3 and H2AK119ub1 maintenance. **A,** Genome browser gene tracks highlighting the location of annotated CpG islands (CGIs) at the indicated genes. **B,** Western Blot showing the levels of MTF2 protein in *MTF2* deleted cell line (*MTF2^-/-^)* with the reporter inserted at the Polycomb target locus*, WT1*. **C,** Western Blot showing the expression levels of *Flag-MTF2* and *Flag-MTF2-EH* overexpressed in *MTF2^-/-^* cells. **D,** Genome browser snapshots of Flag ChIP-seq reads at the reporter locus and the endogenous Polycomb-silenced *FOXQ1* gene in *Flag-MTF2* and *Flag-MTF2-EH* cells after establishment of silencing (+Dox) and 8 days after removal of doxycycline (Dox removal). **E,** Genome browser snapshots of H3K27me3 ChIP-seq reads at the reporter locus and the endogenous Polycomb-silenced *PCDH10* gene in wild-type (WT, *MTF2^+/+^*), *MTF2^-/-^, Flag-MTF2* and *Flag-MTF2-EH* cells after establishment of silencing (+Dox) and 8 days after removal of doxycycline (Dox removal).

**Figure S8.**
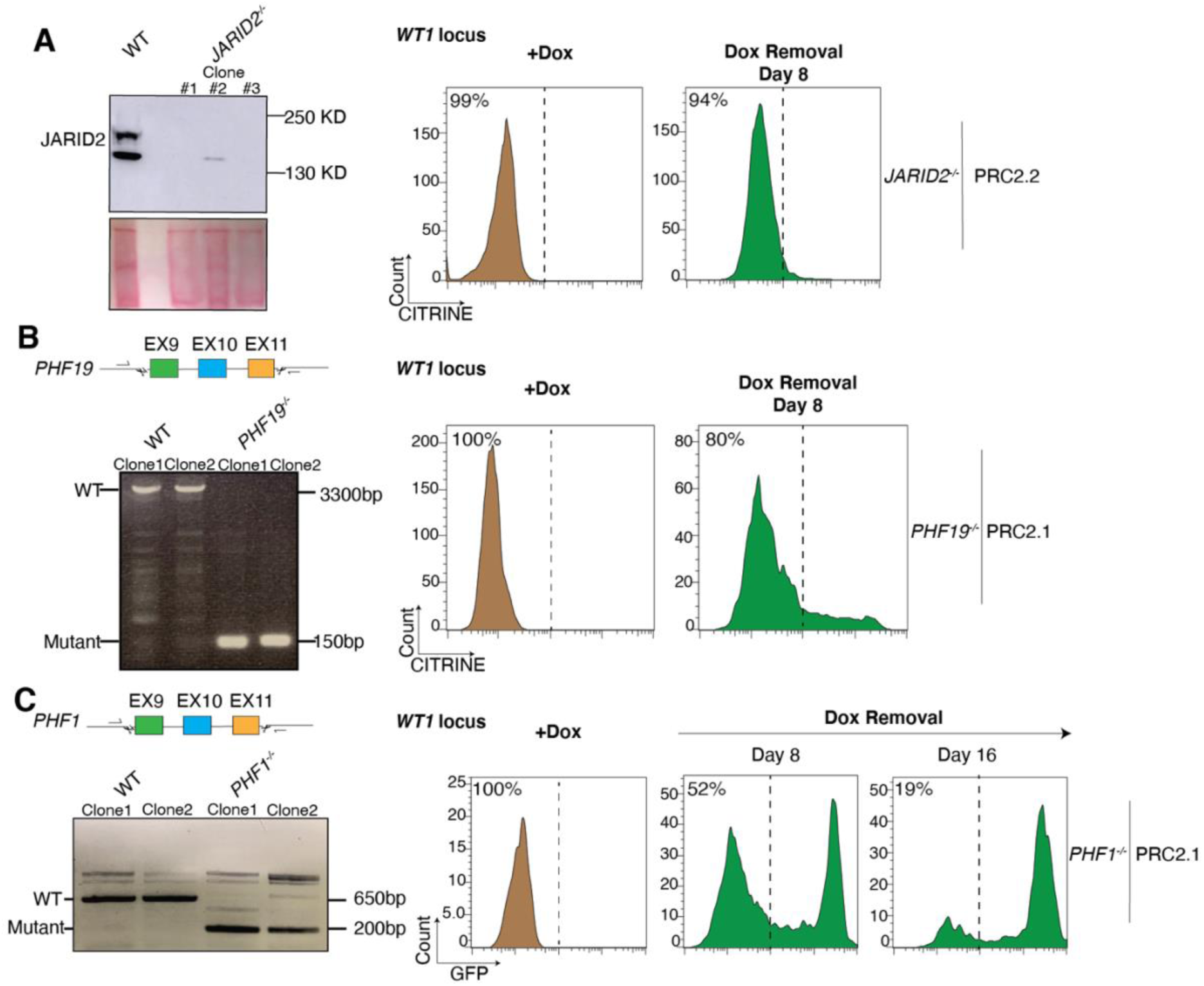
*PHF1* but not *JARID2* or *PHF19,* contribute to epigenetic inheritance of Polycomb silencing. **A,** Western Blot showing the levels of JARID2 protein in *JARID2* deleted cell line (*JARID2^-/-^)* and wild-type cells (left). Flow cytometry histograms showing CITRINE expression after establishment of silencing (+Dox) and 8 days after removal of doxycycline (Dox removal) in *JARID2^-/-^* cells (right). **B,** PCR Genotyping blots validating deletion of exon 9-11 of *PHF19* (left). Flow cytometry histograms showing CITRINE expression after establishment of silencing (+Dox) and 8 days after removal of doxycycline (Dox removal) in *PHF19^-/-^* cells (right). **C,** PCR genotyping blots validating deletion of exon 9-11 of *PHF1* (left). Flow cytometry histograms showing CITRINE expression after establishment of silencing (+Dox) and 8 and 16 days after removal of doxycycline (Dox removal) in *PHF1^-/-^*cells (right).

**Figure S9.**
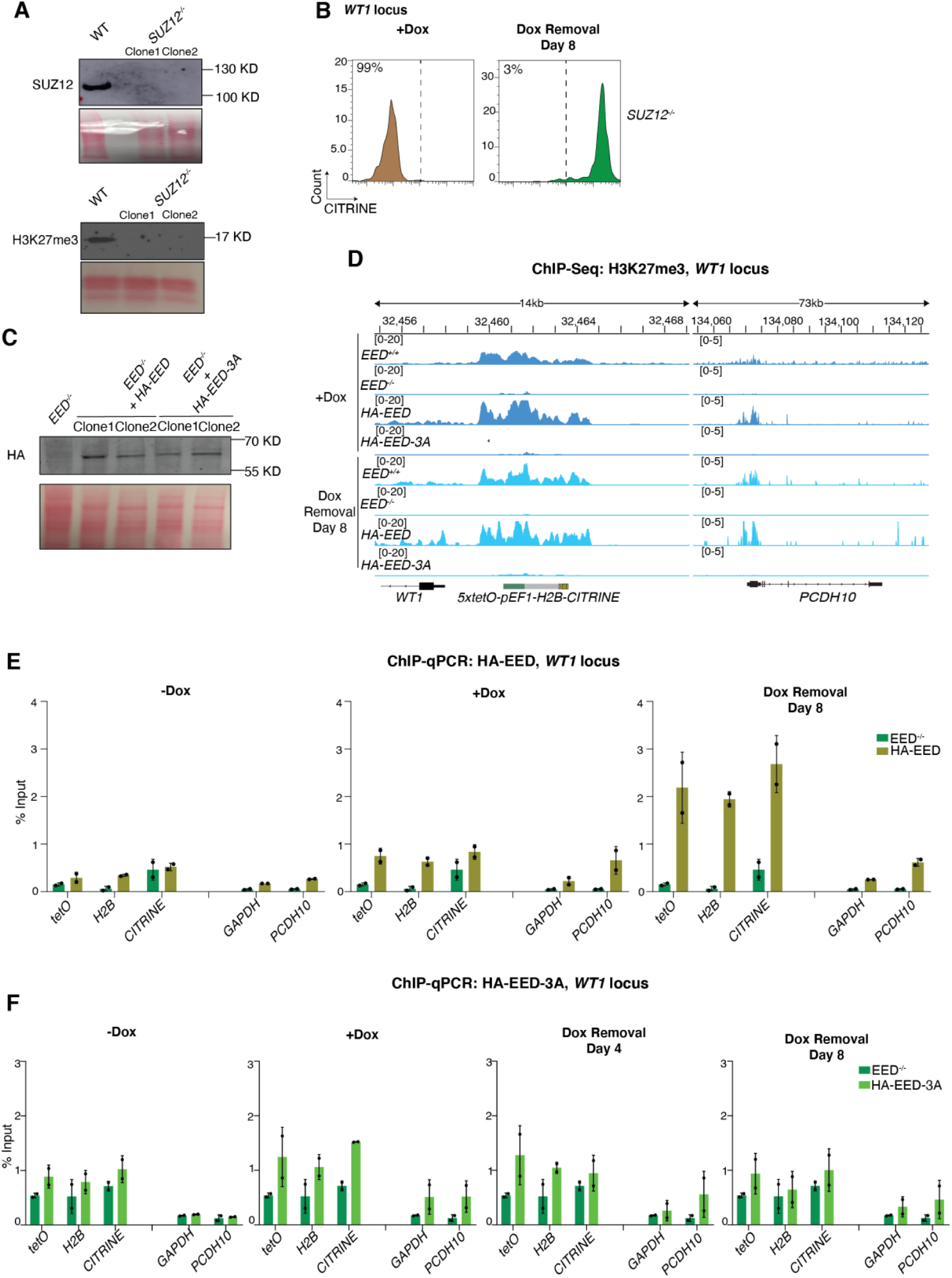
*SUZ12* deletion and the aromatic cage mutant *EED* leads to defects in H3K27me3 catalysis, but the mutant EED can be recruited to chromatin. **A,** Western Blot showing the levels of SUZ12 protein (top) and H3K27me3 (bottom) in *SUZ12* deleted cell line (*SUZ12^-/-^)* and wild-type cells (WT). **B,** Flow cytometry histograms showing CITRINE expression after establishment of silencing (+Dox) and 8 days after removal of doxycycline (Dox removal) in *SUZ12^-/-^* cells. **C,** Western blot showing the expression levels of HA-EED and HA-EED-3A overexpressed in *EED^-/-^*cells. **D,** Genome browser snapshots of H3K27me3 ChIP-seq reads at the reporter locus and the endogenous Polycomb-silenced *PCDH10* gene in *EED^+/+^, EED^-/-^, HA-EED* and *HA-EED-3A* cells after establishment of silencing (+Dox) and 8 days after removal of doxycycline (Dox removal). **E-F,** ChIP-qPCR analysis showing enrichment of HA-EED (**E)** and HA-EED-3A (**F)** expressed in *EED^-/-^* cells, before establishment (-Dox), after establishment (+Dox), and 4 and 8 days after removal of doxycycline (Dox removal) at the reporter locus. *GAPDH* served as a negative control and the native Polycomb-repressed *PCDH10* gene as a positive control. Error bars represent standard deviations.

**Figure S10.**
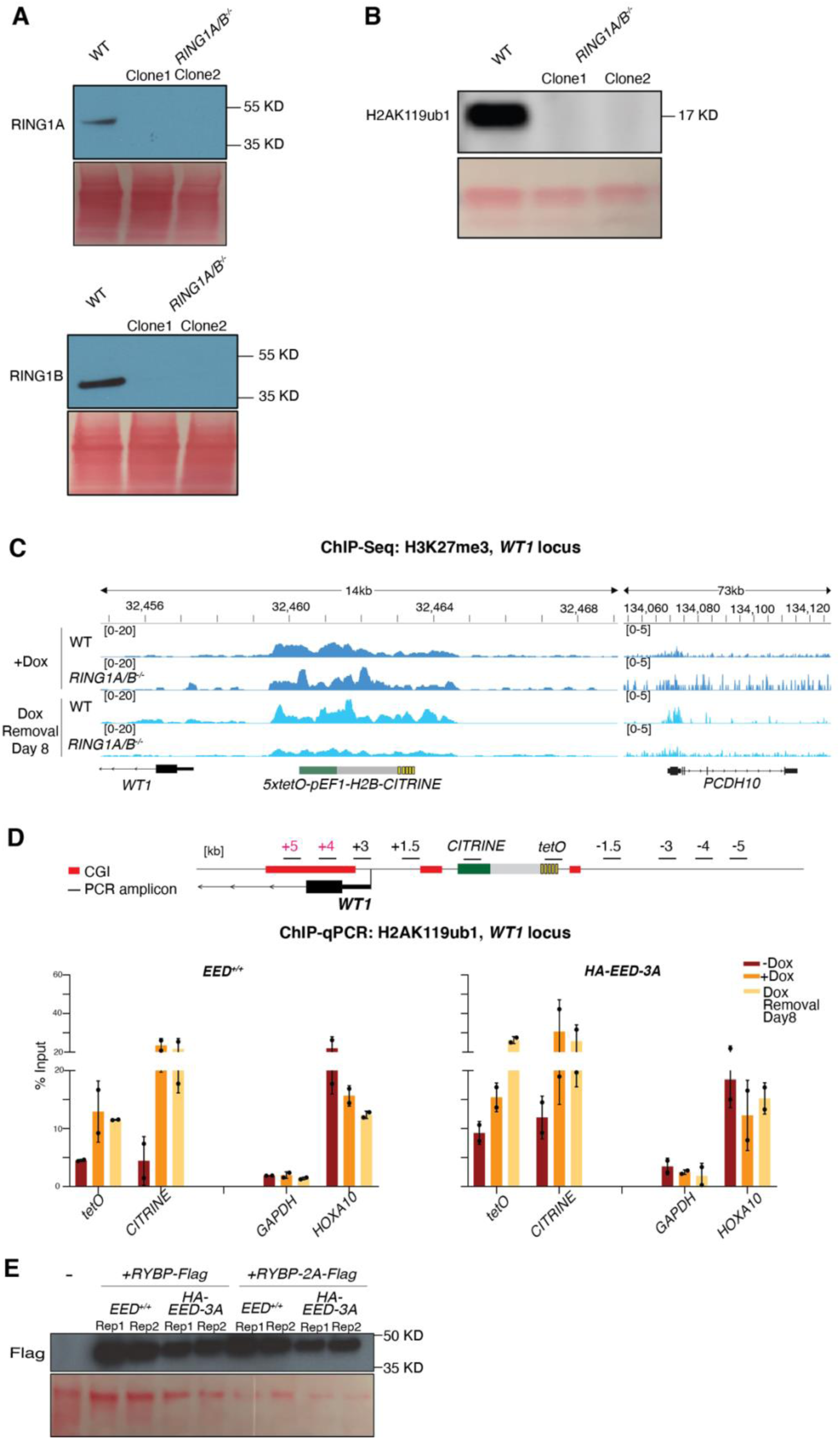
H2AK119ub1 memory depends on RING1A/B and EED but not the ability of EED to bind to H3K27me3. **A,** Western Blot showing the expression levels of RING1A protein (top) and RING1B protein (bottom) in *RING1A/B^-/-^* cells. **B,** Western Blot showing the expression levels of H2AK119ub1 in *RING1A/B^-/-^*cells. **C,** Genome browser snapshots of H3K27me3 ChIP-seq reads at the *WT1* reporter locus and the endogenous Polycomb-silenced *PCDH10* gene in wild-type Control (WT) and *RING1A/B^-/-^* cells after establishment of silencing (+Dox) and 8 days after removal of doxycycline (Dox removal). See Figure 6 for *CITRINE* reporter expression data. **D,** ChIP-qPCR analysis of H2AK119ub1 enrichment at the *CITRINE* reporter, inserted near *WT1*, and surrounding sequences in *EED^+/+^,* and *HA-EED-3A* cells before establishment (-Dox), after establishment (+Dox), and 8 days after removal of doxycycline (Dox removal). *GAPDH* served as a negative control and the native Polycomb-repressed *HOXA10* gene as a positive control. Error bars represent standard deviation. **E,** Western Blot showing the expression levels of wild-type RYBP-Flag and RYBP-2A-Flag proteins in *EED^+/+^* and *HA-EED-3A* cells. (-), non-transfected control cells.

**Table S1.**
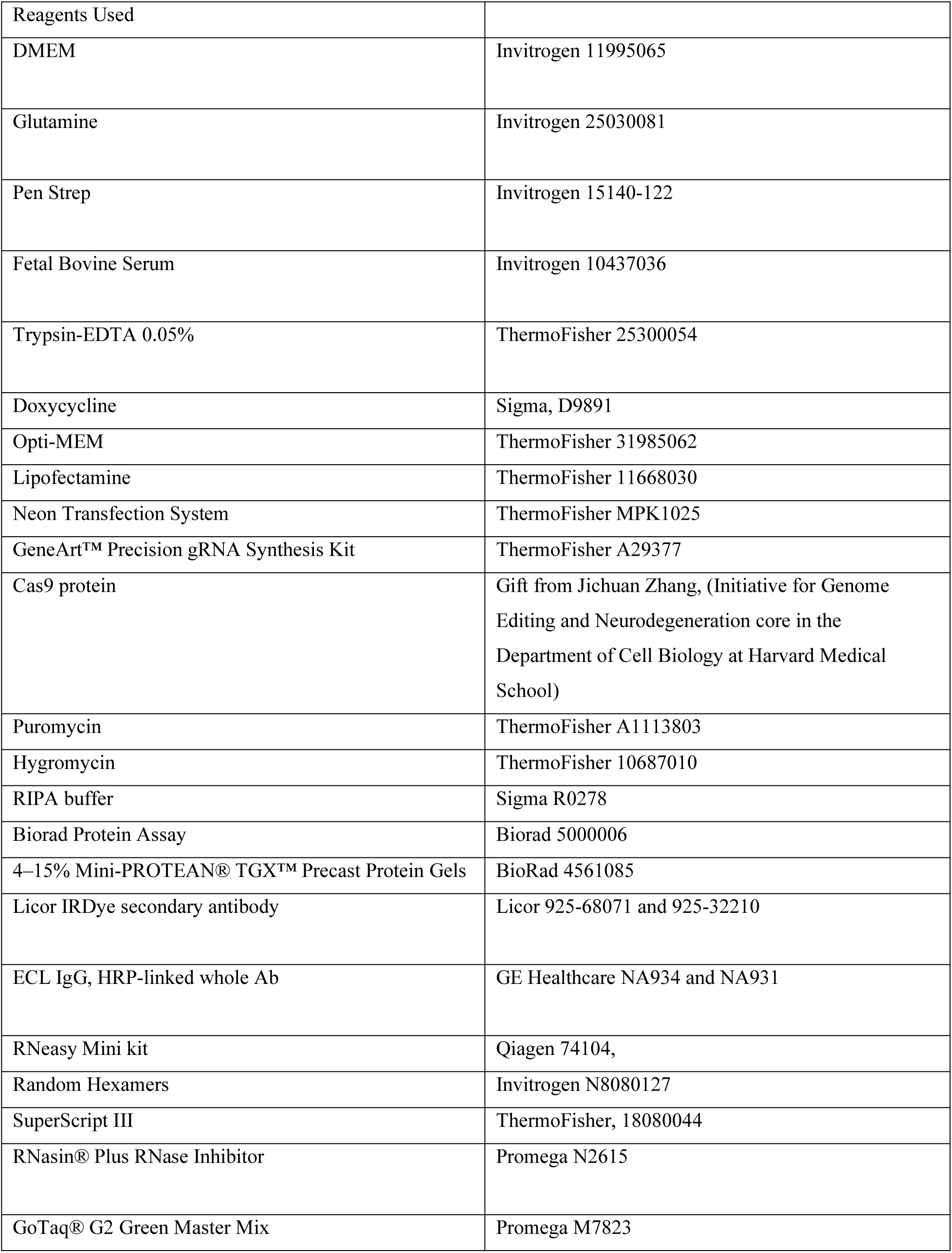
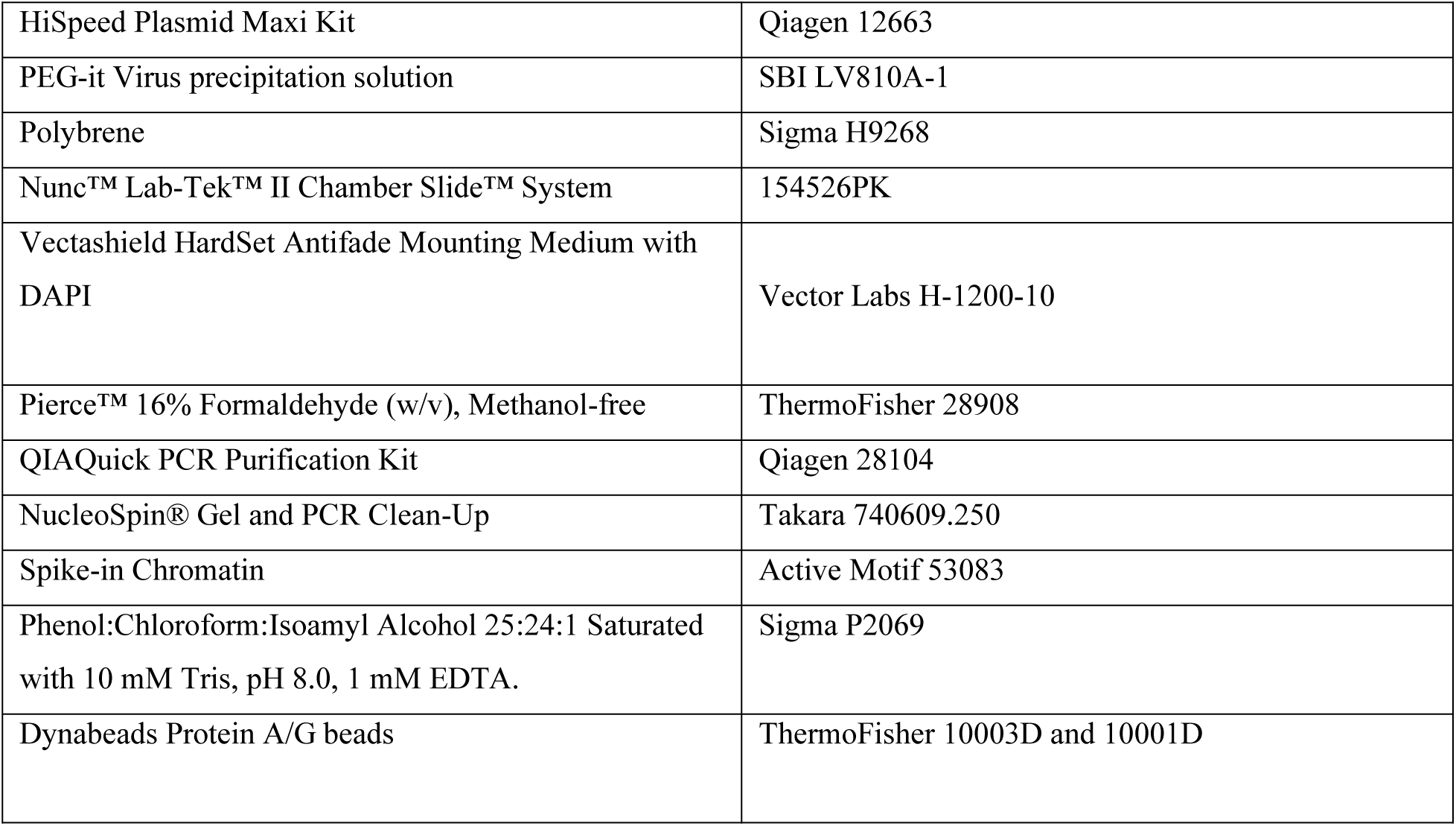
List of reagents used in this study.

| Reagents Used |  |
| --- | --- |
| DMEM | Invitrogen 11995065 |
| Glutamine | Invitrogen 25030081 |
| Pen Strep | Invitrogen 15140-122 |
| Fetal Bovine Serum | Invitrogen 10437036 |
| Trypsin-EDTA 0.05% | ThermoFisher 25300054 |
| Doxycycline | Sigma, D9891 |
| Opti-MEM | ThermoFisher 31985062 |
| Lipofectamine | ThermoFisher 11668030 |
| Neon Transfection System | ThermoFisher MPK1025 |
| GeneArt™ Precision gRNA Synthesis Kit | ThermoFisher A29377 |
| Cas9 protein | Gift from Jichuan Zhang, (Initiative for Genome Editing and Neurodegeneration core in the Department of Cell Biology at Harvard Medical School) |
| Puromycin | ThermoFisher A1113803 |
| Hygromycin | ThermoFisher 10687010 |
| RIPA buffer | Sigma R0278 |
| Biorad Protein Assay | Biorad 5000006 |
| 4–15% Mini-PROTEAN® TGX™ Precast Protein Gels | BioRad 4561085 |
| Licor IRDye secondary antibody | Licor 925-68071 and 925-32210 |
| ECL IgG, HRP-linked whole Ab | GE Healthcare NA934 and NA931 |
| RNeasy Mini kit | Qiagen 74104, |
| Random Hexamers | Invitrogen N8080127 |
| SuperScript III | ThermoFisher, 18080044 |
| RNasin® Plus RNase Inhibitor | Promega N2615 |
| GoTaq® G2 Green Master Mix | Promega M7823 |
| HiSpeed Plasmid Maxi Kit | Qiagen 12663 |
| PEG-it Virus precipitation solution | SBI LV810A-1 |
| Polybrene | Sigma H9268 |
| Nunc™ Lab-Tek™ II Chamber Slide™ System | 154526PK |
| Vectashield HardSet Antifade Mounting Medium with DAPI | Vector Labs H-1200-10 |
| Pierce™ 16% Formaldehyde (w/v), Methanol-free | ThermoFisher 28908 |
| QIAQuick PCR Purification Kit | Qiagen 28104 |
| NucleoSpin® Gel and PCR Clean-Up | Takara 740609.250 |
| Spike-in Chromatin | Active Motif 53083 |
| Phenol:Chloroform:Isoamyl Alcohol 25:24:1 Saturated with 10 mM Tris, pH 8.0, 1 mM EDTA. | Sigma P2069 |
| Dynabeads Protein A/G beads | ThermoFisher 10003D and 10001D |

**Table S2.**
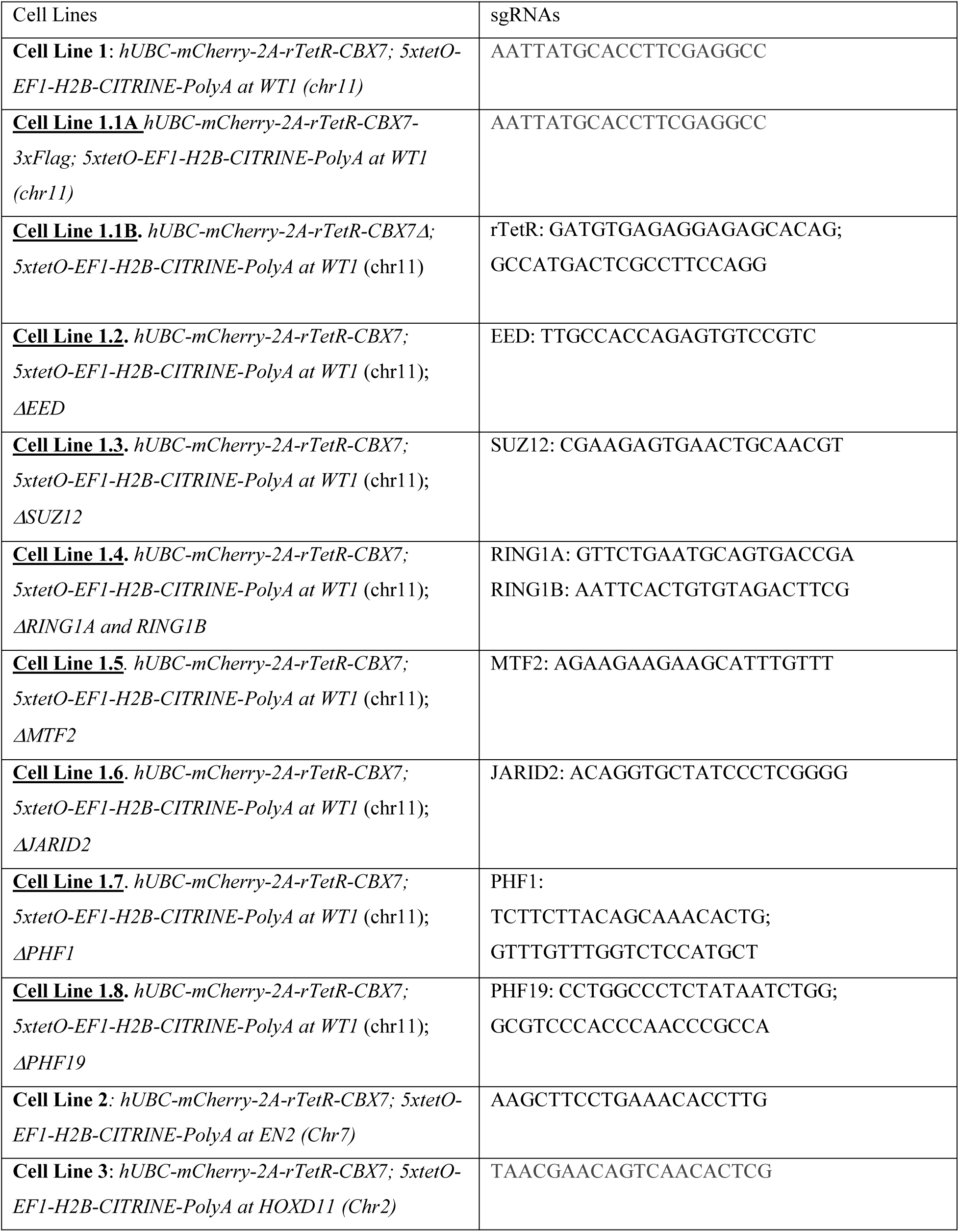
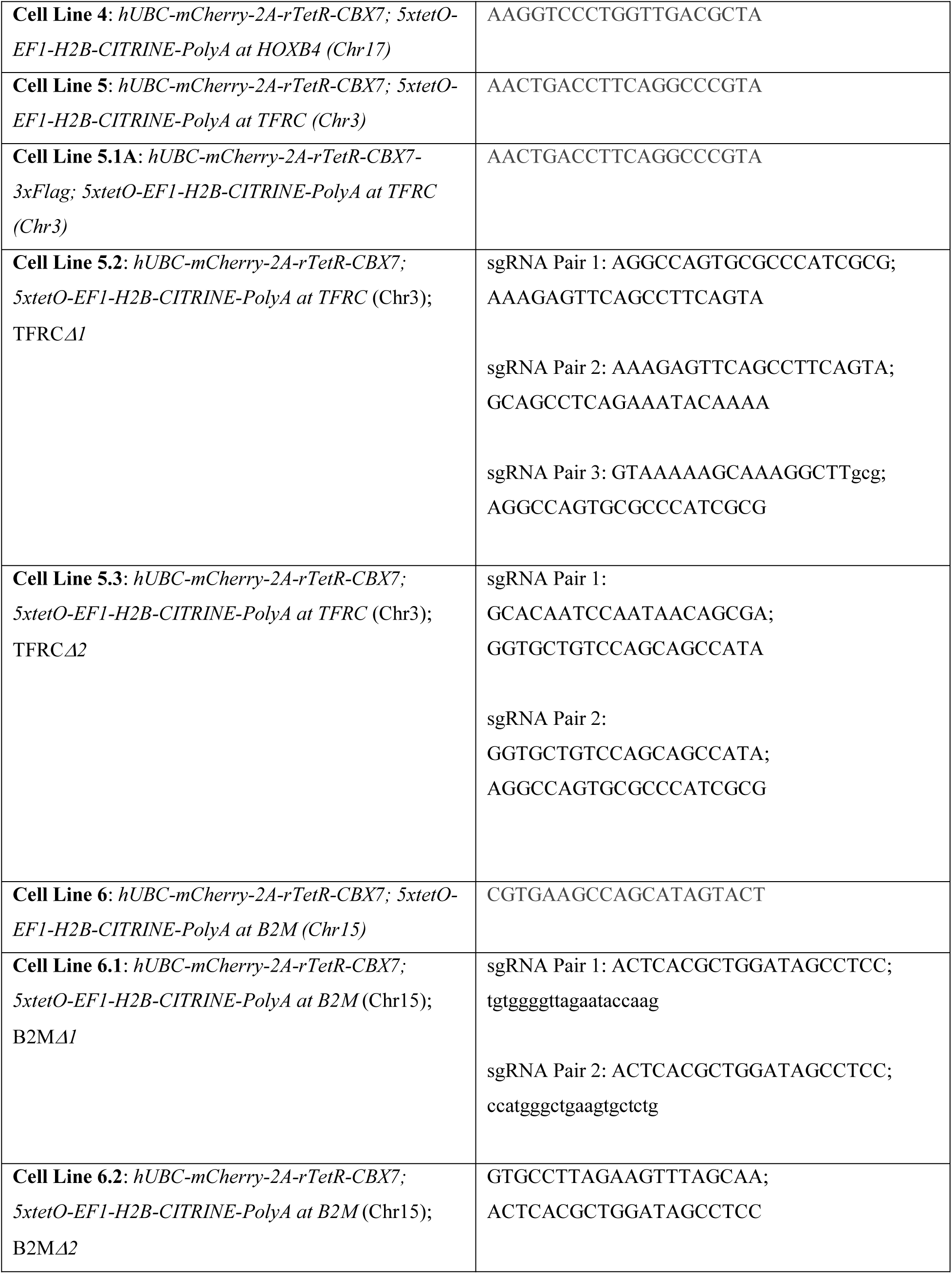
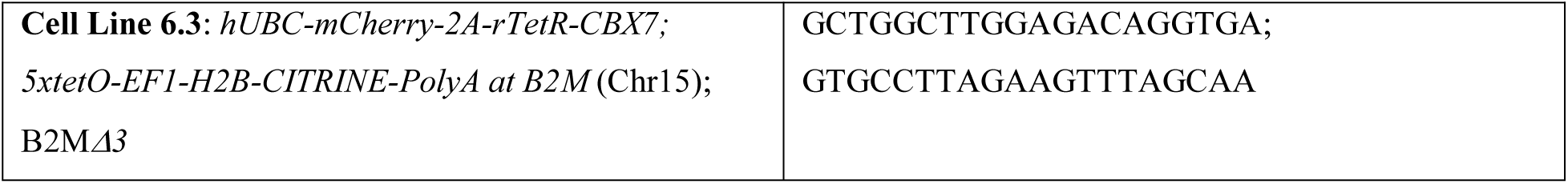
List of the cell lines generated and the sgRNAs used for this study.

**Table S3.**
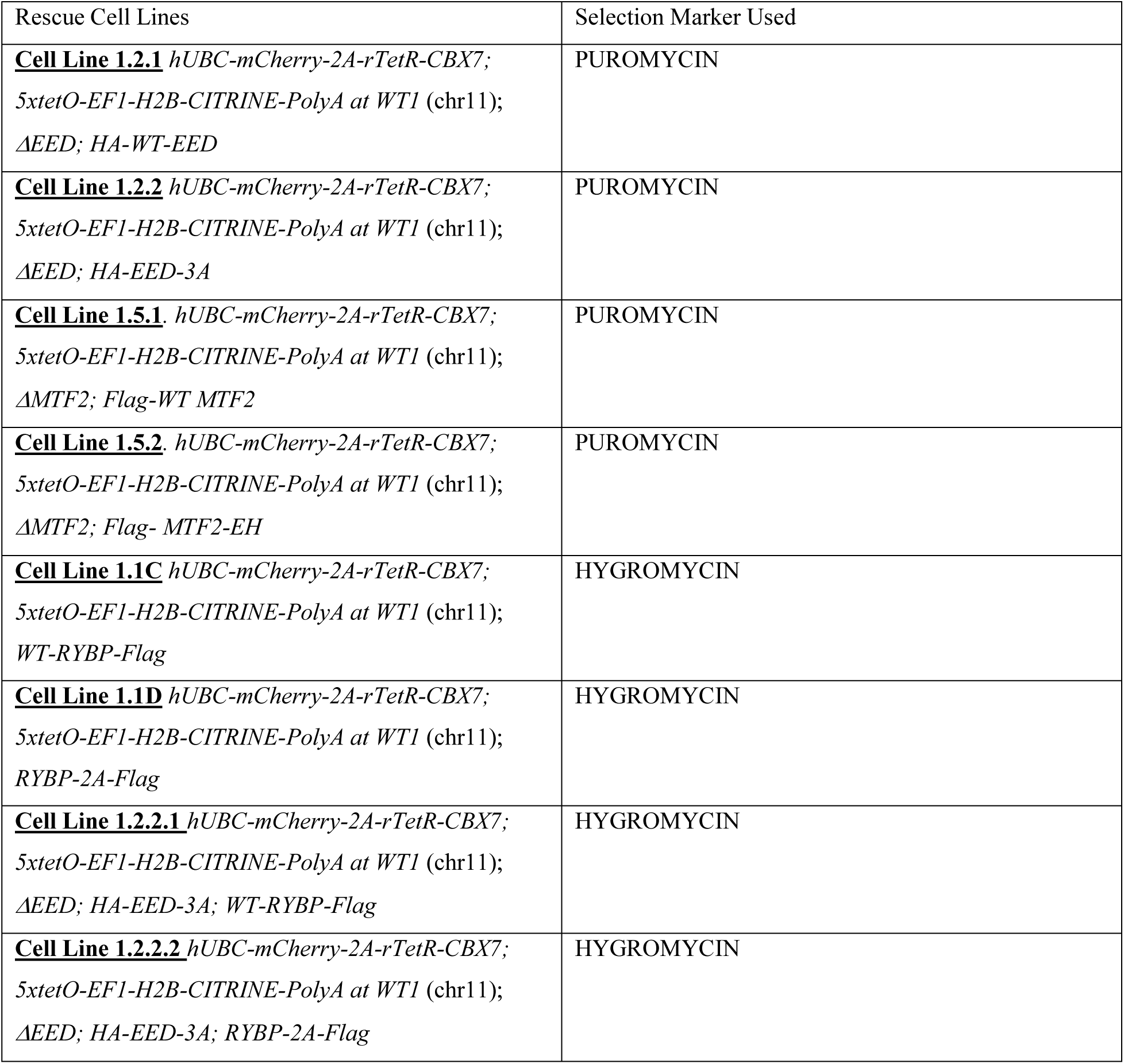
List of the rescue construct cell lines generated and the selection marker used for this study.

| Rescue Cell Lines | Selection Marker Used |
| --- | --- |
| **<u>Cell Line 1.2.1</u>** *hUBC-mCherry-2A-rTetR-CBX7; 5xtetO-EF1-H2B-CITRINE-PolyA at WT1 (chr11); ΔEED; HA-WT-EED* | PUROMYCIN |
| **<u>Cell Line 1.2.2</u>** *hUBC-mCherry-2A-rTetR-CBX7; 5xtetO-EF1-H2B-CITRINE-PolyA at WT1 (chr11); ΔEED; HA-EED-3A* | PUROMYCIN |
| **<u>Cell Line 1.5.1</u>** *hUBC-mCherry-2A-rTetR-CBX7; 5xtetO-EF1-H2B-CITRINE-PolyA at WT1 (chr11); ΔMTF2; Flag-WT MTF2* | PUROMYCIN |
| **<u>Cell Line 1.5.2</u>** *hUBC-mCherry-2A-rTetR-CBX7; 5xtetO-EF1-H2B-CITRINE-PolyA at WT1 (chr11); ΔMTF2; Flag- MTF2-EH* | PUROMYCIN |
| **<u>Cell Line 1.1C</u>** *hUBC-mCherry-2A-rTetR-CBX7; 5xtetO-EF1-H2B-CITRINE-PolyA at WT1 (chr11); WT-RYBP-Flag* | HYGROMYCIN |
| **<u>Cell Line 1.1D</u>** *hUBC-mCherry-2A-rTetR-CBX7; 5xtetO-EF1-H2B-CITRINE-PolyA at WT1 (chr11); RYBP-2A-Flag* | HYGROMYCIN |
| **<u>Cell Line 1.2.2.1</u>** *hUBC-mCherry-2A-rTetR-CBX7; 5xtetO-EF1-H2B-CITRINE-PolyA at WT1 (chr11); ΔEED; HA-EED-3A; WT-RYBP-Flag* | HYGROMYCIN |
| **<u>Cell Line 1.2.2.2</u>** *hUBC-mCherry-2A-rTetR-CBX7; 5xtetO-EF1-H2B-CITRINE-PolyA at WT1 (chr11); ΔEED; HA-EED-3A; RYBP-2A-Flag* | HYGROMYCIN |

**Table S4.**
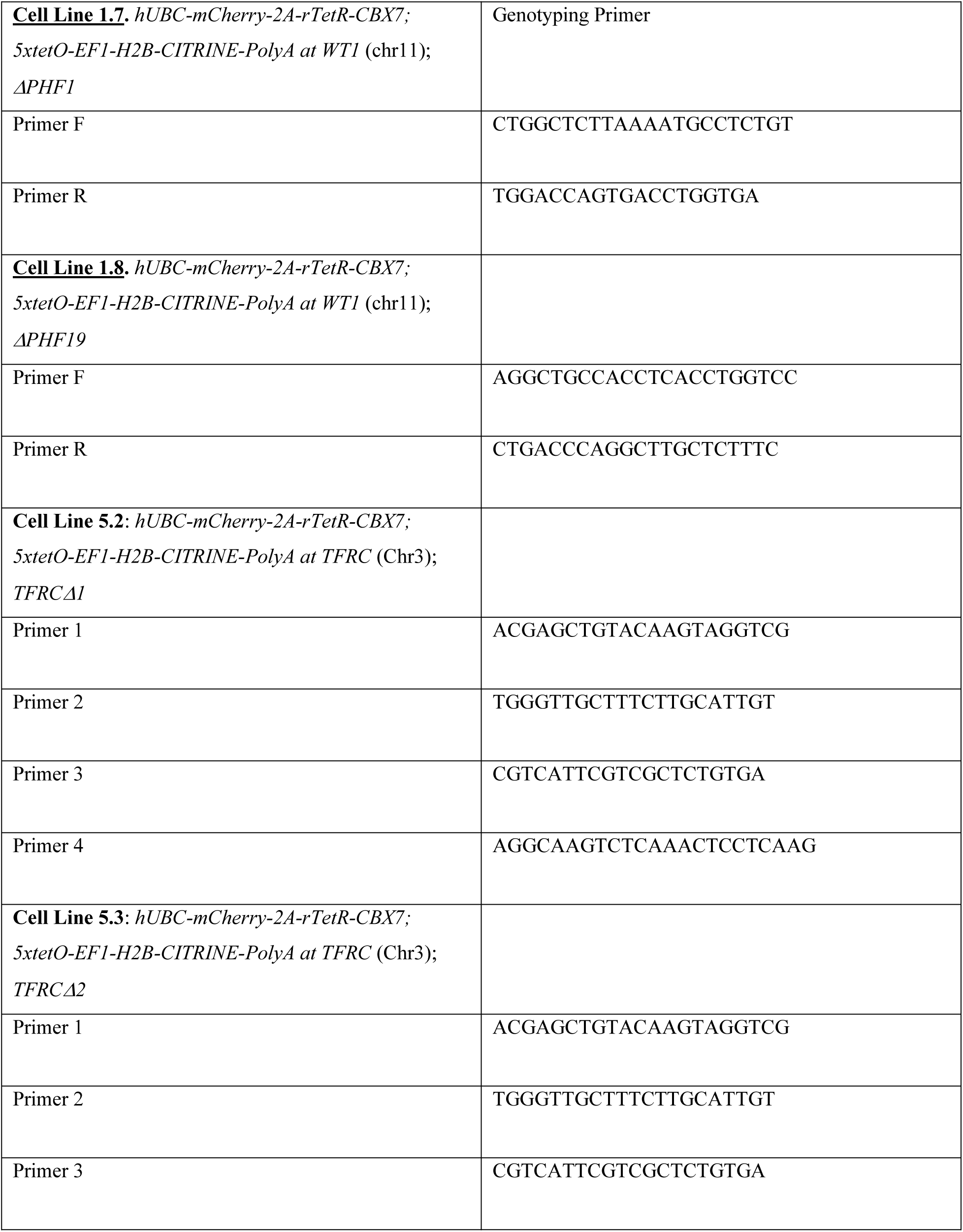
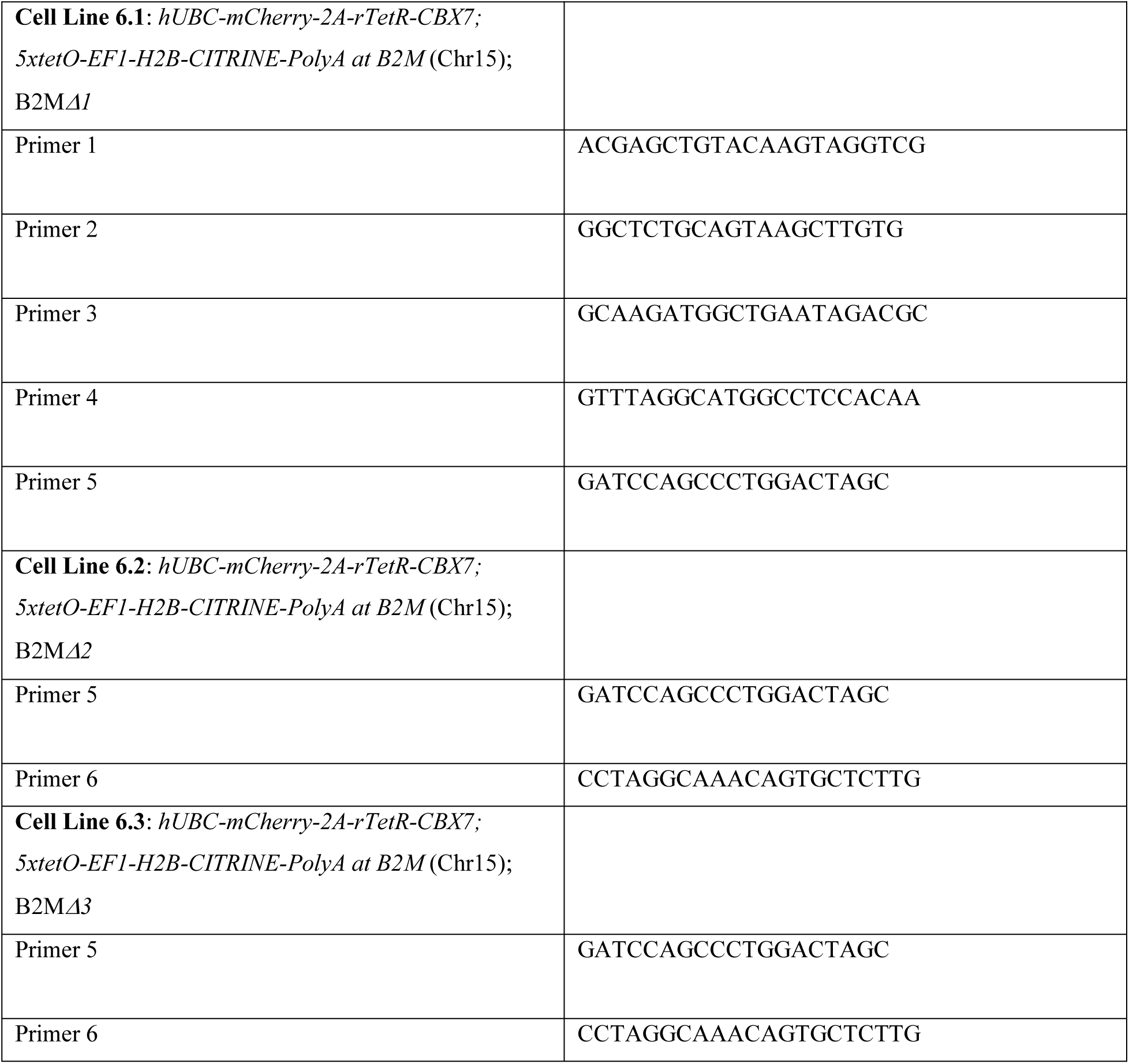
List of Genotyping Primers used in this study.

**Table S5.** List of Antibodies used in this study.

| Antibodies | Source | Cat# | Application |
| --- | --- | --- | --- |
| Anti-RING1B | Cell Signaling Technology | 5694S | WB, 1:200 |
| Anti-RING1A | Cell Signaling Technology | 13069S | WB, 1:200 |
| Anti-BETA ACTIN | Abcam | mAbcam8224 | WB, 1:1000 |
| Anti-FLAG | Sigma | F3165 | WB, 1:5000<br>ChIP, 4ug |
| Anti-SUZ12 | Cell Signaling Technology | 3737 | IB, 1:1000 |
| Anti-H3K27me3 | Cell Signaling Technology | 9733 | WB, 1:1000<br>ChIP, 3ug |
| Anti-H2AK119ub1 | Cell Signaling Technology | 8240T | WB, 1:1000<br>ChIP, 2ug |
| Anti-MTF2 | ProteinTech | 16208-1-AP | WB, 1:100<br>ChIP, 4ug |
| Anti-JARID2 | Novus Biologics | NB100-2214SS | WB, 1:500 |
| Anti-HA | ThermoFisher | 26183-HRP | WB, 1:2000 |
| Anti-HA Tag Monoclonal Antibody (2-2.2.14) | Thermo-Fisher | 26183 | ChIP, 2ug |
| Anti-CBX7 | Abcam | ab21873 | WB, 1:1000 |
| Anti-Histone H2Av antibody | Active Motif | 61686 | ChIP-Seq Spike in: 2ug |
| Anti-B2M | Cell Signaling | 12851 | WB, 1:200 |
| Anti- TFRC | Cell Signaling | 13113T | WB, 1:200 |
| Anti-RYBP | Sigma | PRS2227 | WB, 1:200 |

**Table S6.**
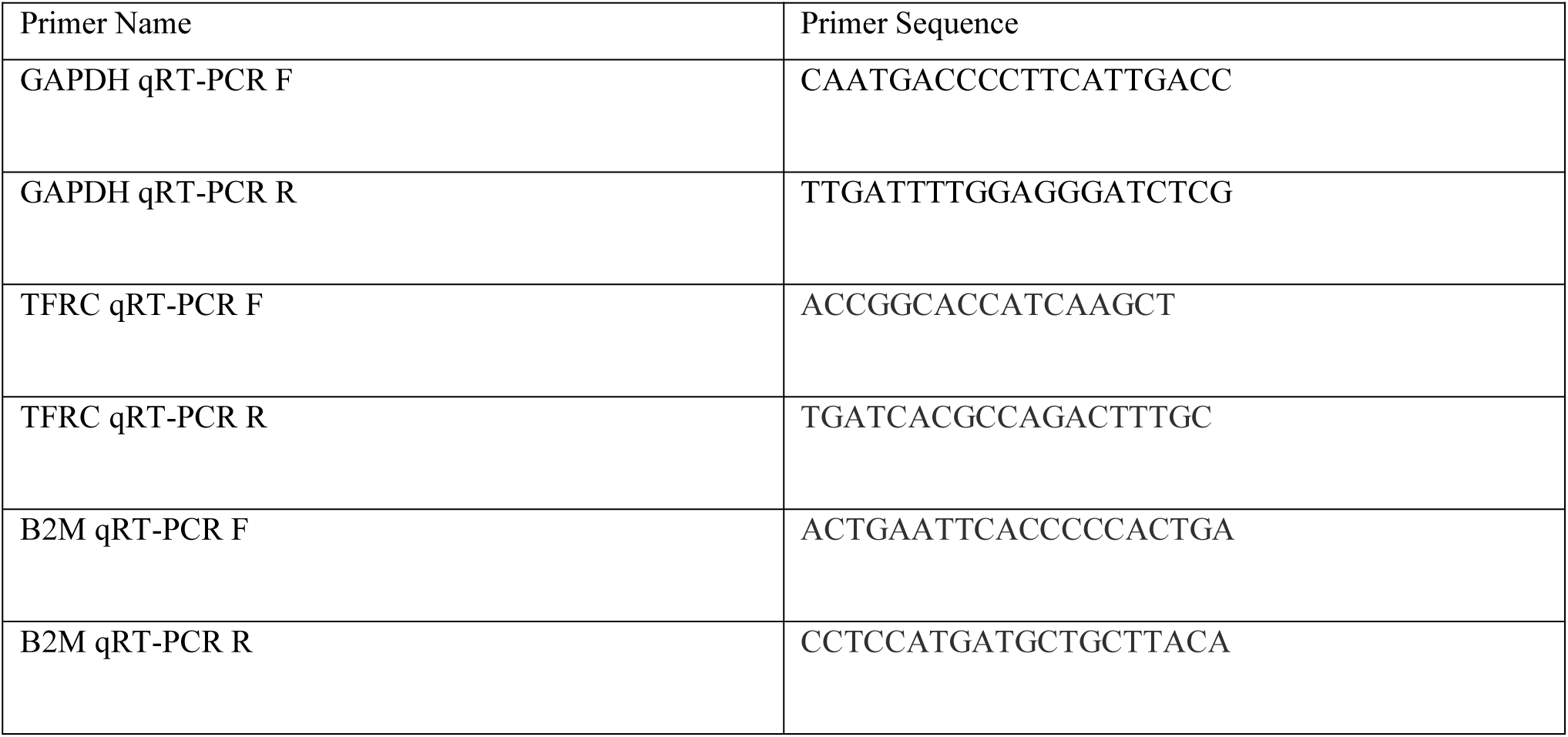
qRT-PCR primers used in this study.

**Table S7.**
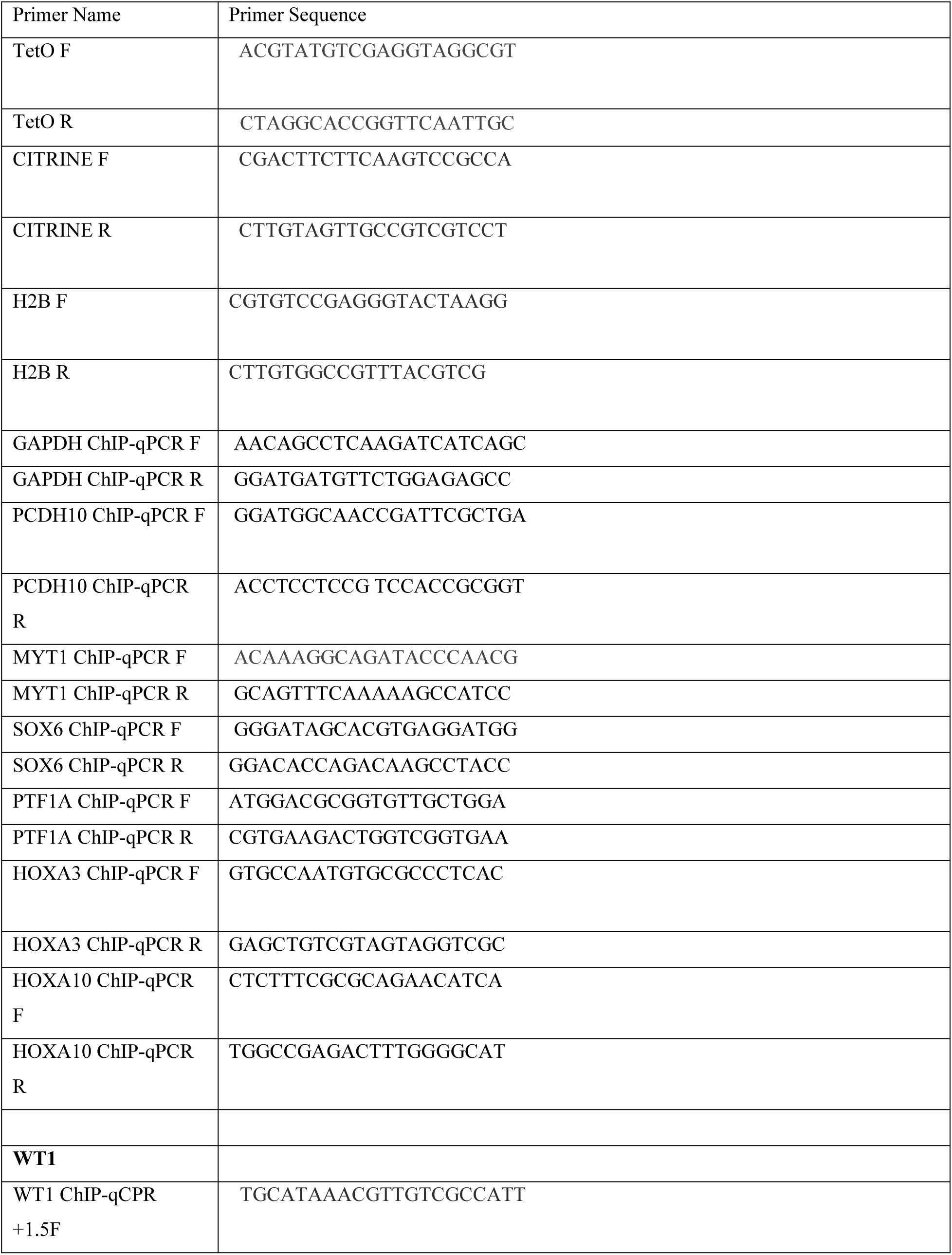
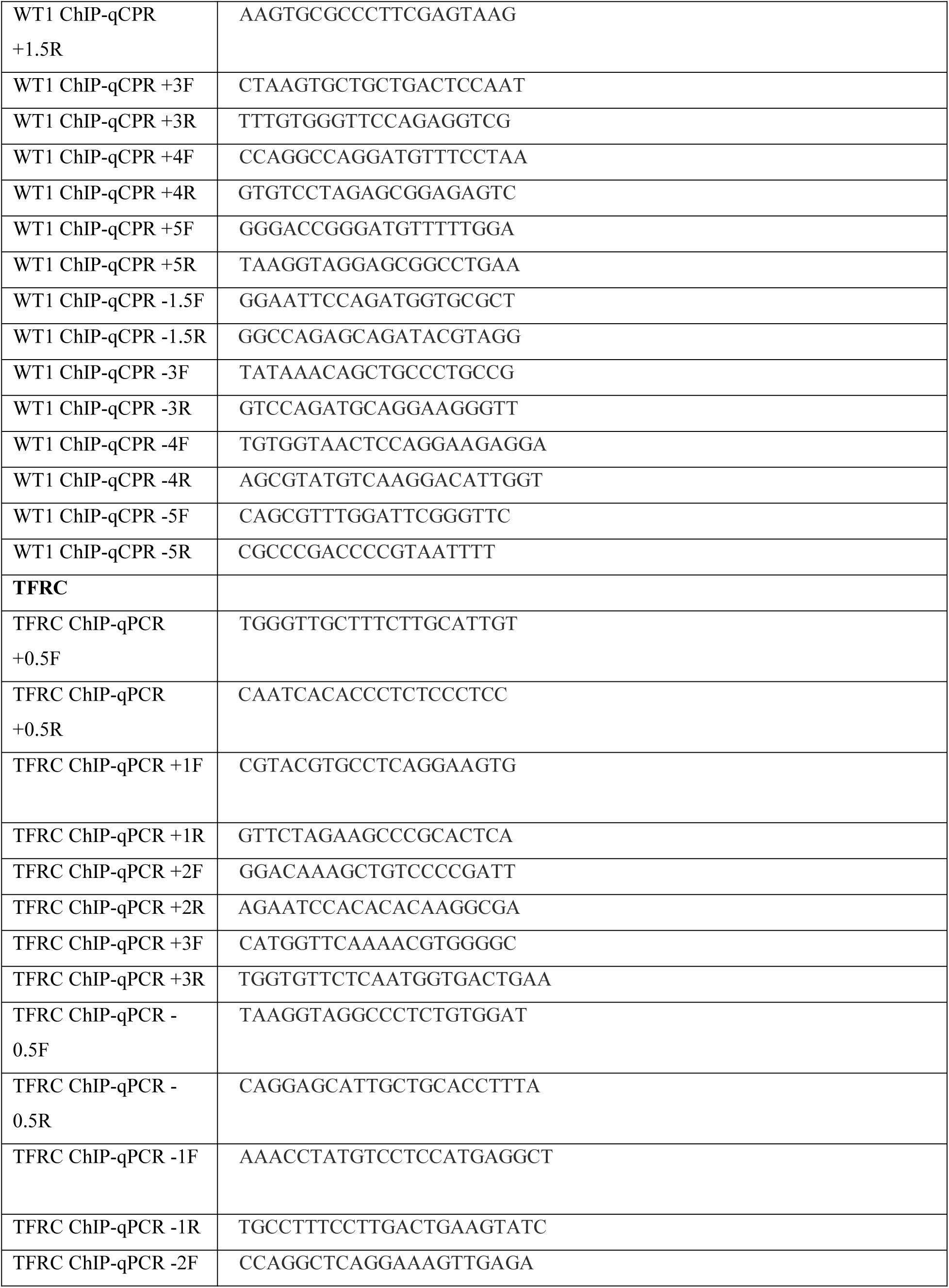
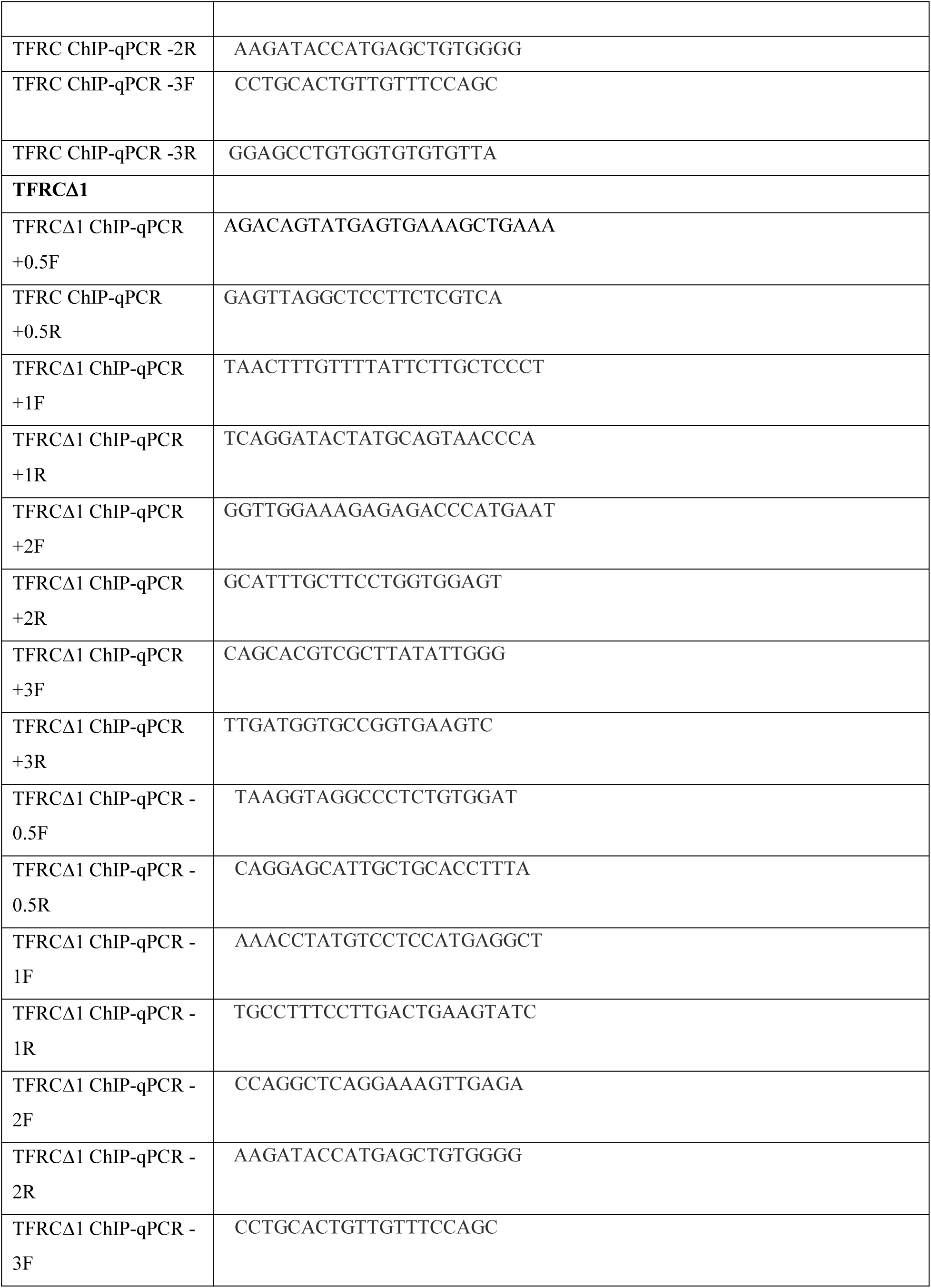
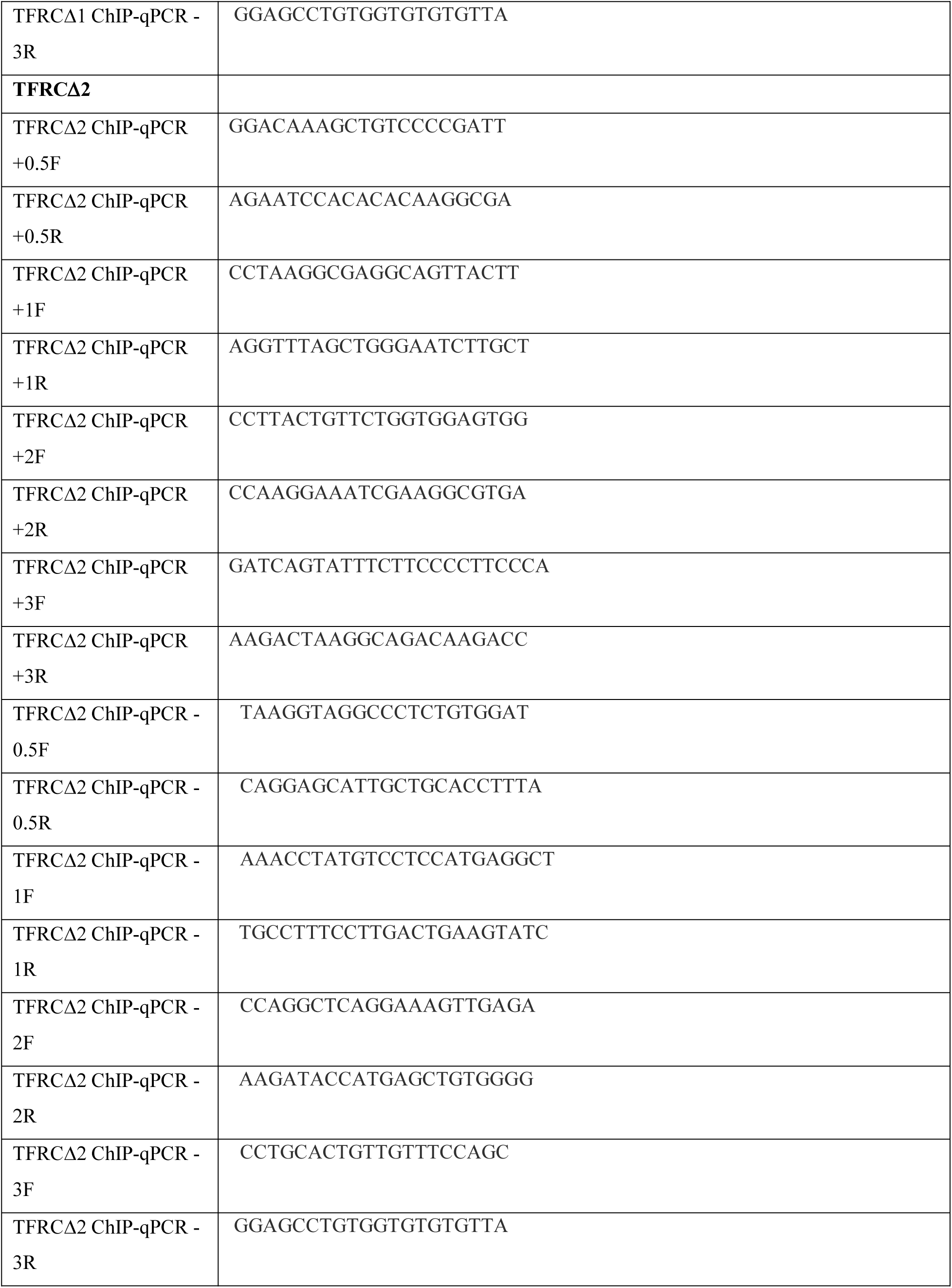
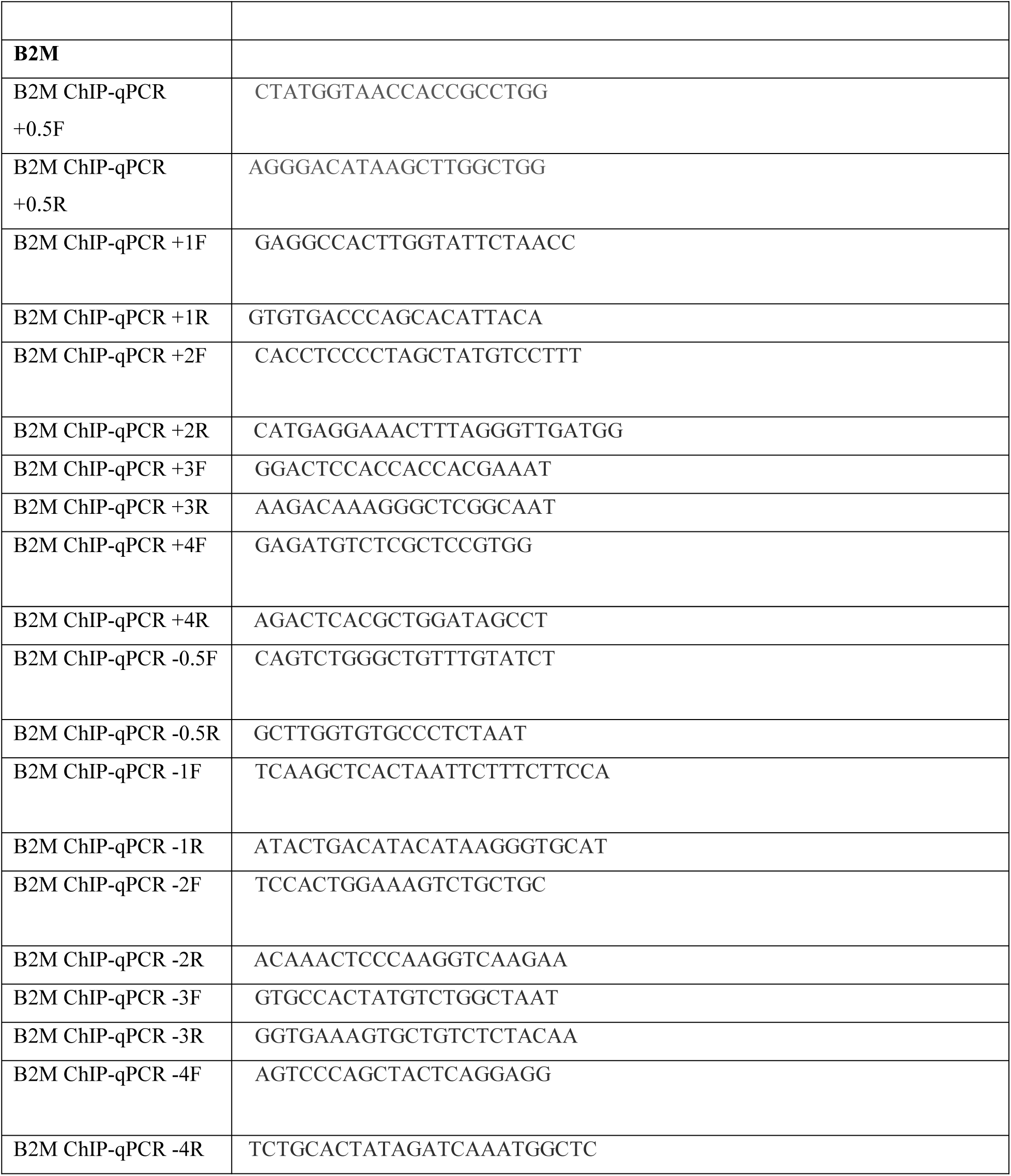
ChIP-qPCR primers used in this study.

**Table S8.** List of data sources from public databases.

| **File Name** | **GEO Accession #/Source Information** |
| --- | --- |
| hESCs H3K27me3 ChIP-Seq | GSM1185386 |
| hESCs RNA-seq | GSM672836 |
| HEK293FT H3K27me3 ChIP-Seq | GSM4239945 |
| HEK293FT H2AK119ub1 ChIP-Seq | GSM4502558 |
| HEK293FT RNA-seq | GSM5343713 |
| cMYC ChIP | GSM3360524 |
| HEK293 Bisulfite Seq | GSM683769 |
| CGIs | UCSC Genome browser |
| Human Protein Atlas | proteinatlas.org |
| Transcription Factor ChIP | UCSC Genome browser |
| MTF2 PDB file | 10.2210/pdb5XFR/pdb (Li et al, 2017)<sup>36</sup> |
| EED PDB file | 10.2210/pdb3IIW/pdb (Margueron et al, 2009)<sup>21</sup> |

**Table S9.** List of Softwares used in this study.

|  |
| --- |
| bowtie2(v2.2.9) |
| deeptool (v3.0.2) |
| Samtool (v1.3.1) |
| Integrative Genomics Viewer (v2.12.0) |
| BD Biosciences-FACsDIVA |
| BD Biosciences CellQuestPro |
| FlowJo (v10.5.3) |
| NIS-Elements imaging software |
| ImageJ (v1.0) |
| GraphPad Prism 7.0c |
| https://benchling.com |
| https://chopchop.cbu.uib.no/ |

